# Formation and Retrieval of Cell Assemblies in a Biologically Realistic Spiking Neural Network Model of Area CA3 in the Mouse Hippocampus

**DOI:** 10.1101/2024.03.27.586909

**Authors:** Jeffrey D. Kopsick, Joseph A. Kilgore, Gina C. Adam, Giorgio A. Ascoli

## Abstract

The hippocampal formation is critical for episodic memory, with area Cornu Ammonis 3 (CA3) a necessary substrate for auto-associative pattern completion. Recent theoretical and experimental evidence suggests that the formation and retrieval of cell assemblies enable these functions. Yet, how cell assemblies are formed and retrieved in a full-scale spiking neural network (SNN) of CA3 that incorporates the observed diversity of neurons and connections within this circuit is not well understood. Here, we demonstrate that a data-driven SNN model quantitatively reflecting the neuron type-specific population sizes, intrinsic electrophysiology, connectivity statistics, synaptic signaling, and long-term plasticity of the mouse CA3 is capable of robust auto-association and pattern completion via cell assemblies. Our results show that a broad range of assembly sizes could successfully and systematically retrieve patterns from heavily incomplete or corrupted cues after a limited number of presentations. Furthermore, performance was robust with respect to partial overlap of assemblies through shared cells, substantially enhancing memory capacity. These novel findings provide computational evidence that the specific biological properties of the CA3 circuit produce an effective neural substrate for associative learning in the mammalian brain.

## Introduction

Episodic memory is a fundamental cognitive operation that links together the contents of a present experience – spatial, temporal, sensory, and emotional – for future recall [1–4]. The hippocampal formation (HPF) is a critical substrate for episodic memory formation and retrieval, with area Cornu Ammonis 3 (CA3) crucial for auto-associative memories [5]. Auto-association and pattern completion are two circuit functions that involve the storage of individual experiences and their recall from a partial cue, respectively [6]. Neurophysiological studies highlight that these experiences are represented by the concurrent firing of a group or groups of excitatory pyramidal cells (PCs), known as neuronal ensembles or cell assemblies [7,8]. Additionally, empirical evidence reveals a synaptic basis for these experiences, where the order and timing of spikes via long-term spike timing-dependent plasticity (STDP) is a key factor in strengthening synaptic conductance at PC-PC synapses [9].

Open questions stemming from CA3 as a substrate for memory regard the quality of experience remembered, and the number of stored experiences: how well does CA3 recall experiences, and what is the memory capacity of CA3? For the first, the quality may be based on how the learned experience is encoded by cell assemblies and their corresponding connections, where changes in the amplitude of excitatory postsynaptic potentials [10] and number of AMPA receptors at the terminals of postsynaptic PCs can occur [9,11–13]. Additionally, the amount of information provided in the form of a cue to these cells can lead to re-activation of the memory through pattern completion [14]. These mechanisms also depend not only on the specific input-output properties of CA3 PCs [15–17], but also on considerably diverse inhibitory interneurons [18].

For the second, theoretical and empirical evidence suggests that there are four key factors in determining the capacity for memories in CA3: the number of PCs, the probability of connection between PCs, the size of cell assemblies, and the amount of overlap between cell assemblies. Estimates for the number of neurons, the PC-PC connection probability, and the size of cell assemblies have been offered based on various assumptions [19,20]. Additionally, estimates have been provided for the percentage of cells shared between cell assemblies, and the shared cells between assemblies provide a neural substrate for associations that enable representations of specific episodic memories [21,22]. Estimates for the memory capacity of CA3 have been offered based on these factors in rats, though, to our knowledge, not in mice. However, these estimates relied on network models that did not reflect the neural and connection type diversity of the CA3 circuit [19,20,23].

Hippocampome.org is an open access knowledge base of distinct *neuron types* in the rodent HPF [24,25]. This resource identifies neuron types based on their primary neurotransmitter (glutamate or GABA) and the presence of axons and dendrites across distinct layers of each cytoarchitectonic area of the HPF: entorhinal cortex, dentate gyrus, CA3, CA2, CA1, and subiculum. Hippocampome.org provides for each neuron type experimental data regarding the expression of specific molecules [26], biophysical membrane properties [27], electrophysiological firing patterns *in vitro* and *in vivo* [28,29] and population size [30,31]. Additionally, Hippocampome.org quantifies the connection probability and synaptic signals of directional pairs formed between a pre- and post-synaptic neuron type, known as *potential connections*, which are based on their axonal and dendritic distributions [32–35]. Also available on this web portal are computational models of neuronal excitability [36] and short-term synaptic plasticity [37] using the Izhikevich and Tsodyks-Markram formalisms, respectively.

Utilizing Hippocampome.org, we previously created a computational circuit model of the mouse CA3 that featured a selection of neuron types and potential connections chosen to represent the neural diversity of this area [38]. Additionally, the *in silico* implementation of this model as a spiking neural network (SNN) in the GPU-based simulation environment CARLsim6 can capture the individual spike times of every neuron, and can track changes in synaptic weight at each connection [39]. This makes the Hippocampome derived CA3 SNN particularly useful for elucidating mechanisms for auto-association and pattern completion.

The present work investigates whether a SNN that reflects the scale, diversity, and biological properties of the mouse CA3 can form and retrieve patterns via cell assemblies. We demonstrate that this SNN has activity consistent with what has been observed *in vivo*, and that patterns are auto-associated and completed robustly with minimally informative cues that stem from cell assembly formation and retrieval, respectively. Additionally, we report that a range of assembly sizes can support pattern completion after a limited number of repeated presentations. Furthermore, when cells are shared between assemblies, auto-association and pattern completion remain nearly unaltered, suggesting that individual representations can be strongly retrieved while still providing a basis for overlapping experiences. Moreover, this finding offers a potential mechanism supporting a substantial expansion of memory capacity in the CA3 circuit.

## Results

### Can a full-scale CA3 SNN store and retrieve patterns via cell assemblies?

To answer this first research question, we utilize our full-scale SNN of the mouse CA3 [38], which exhibited rhythmic network activity that was stable and robust in response to synchronous or asynchronous transient inputs, reflecting resting-state behaviors [38]. This model consisted of 8 neuron types and 51 connection types and was instantiated with 84,053 neurons and 176 million connections (Figure 1A; Tables 1 and 2). Starting from this architecture, we sought to understand how CA3 could embed experiences occurring during wakefulness via cell assemblies for later recall. To create cell assemblies, a symmetric STDP learning rule was implemented in the SNN [12]: Δ*w* = *Ae*^−|Δ*t*|/*τ*^, where *A* determines the peak amplitude of weight change, *τ* is the decay time constant, and Δt is the time difference between the post- and pre-synaptic spikes. Values for each parameter were set to best approximate the symmetric exponential decay curve observed experimentally [12] (Materials and Methods; Figure 1B).

**Figure 1:**
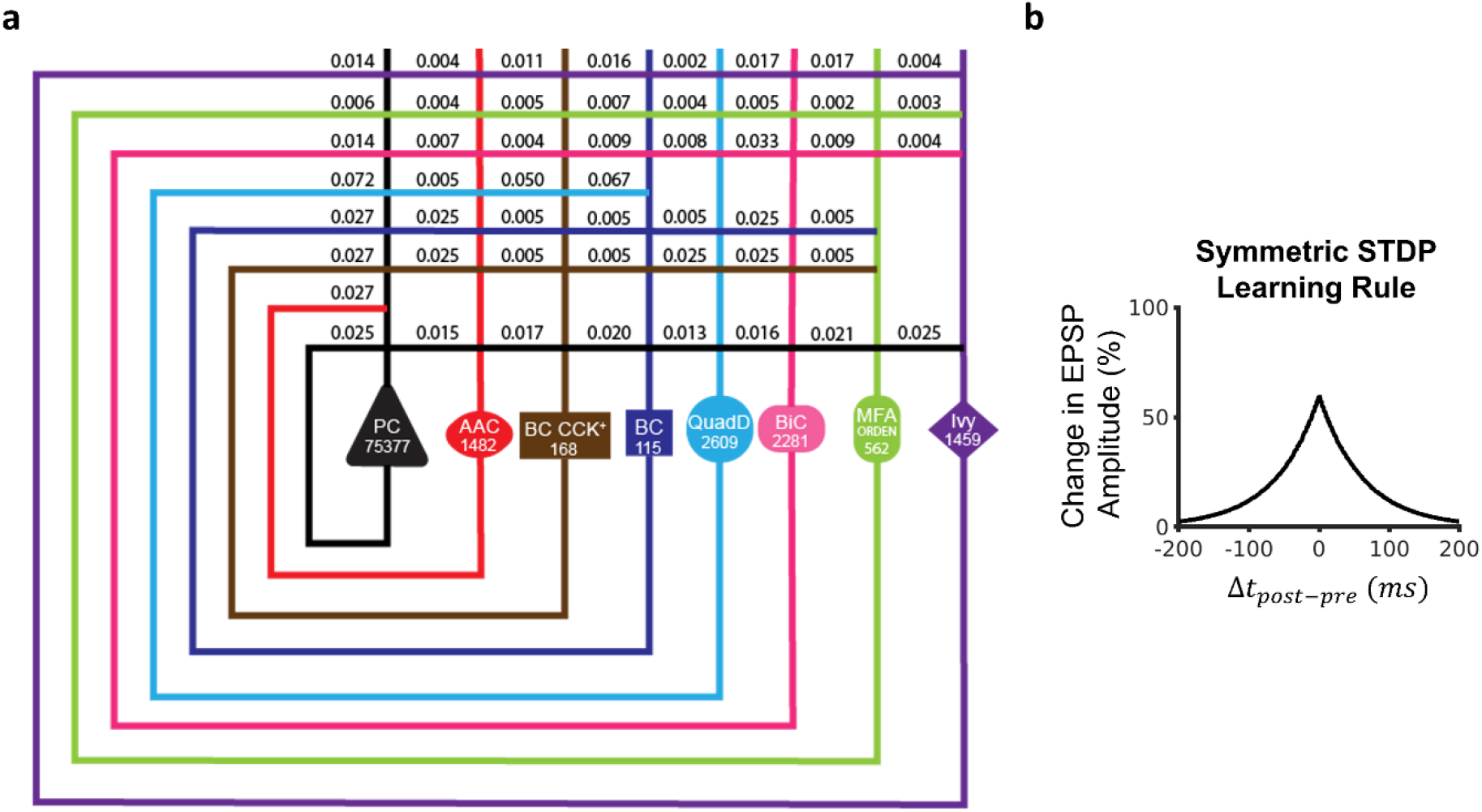
Full-scale CA3 SNN with long-term excitatory synaptic plasticity. (a) Circuit schematic of the CA3 SNN. Cell counts for each neuron type are displayed in the corresponding soma symbol, and probabilities of connection between pairs of neuron types are listed at points of axonal-dendritic overlap. (b) A broad symmetric STDP window promotes synaptic potentiation between concomitantly firing Pyramidal cells, reflecting each pattern. PC = Pyramidal cell; AAC = Axo-axonic cell; BC CCK+ = Basket CCK+; BC = Basket cell; QuadD = QuadD-LM; BiC = Bistratified cell; MFA ORDEN = Mossy Fiber-Associated ORDEN.

**Table 1.**
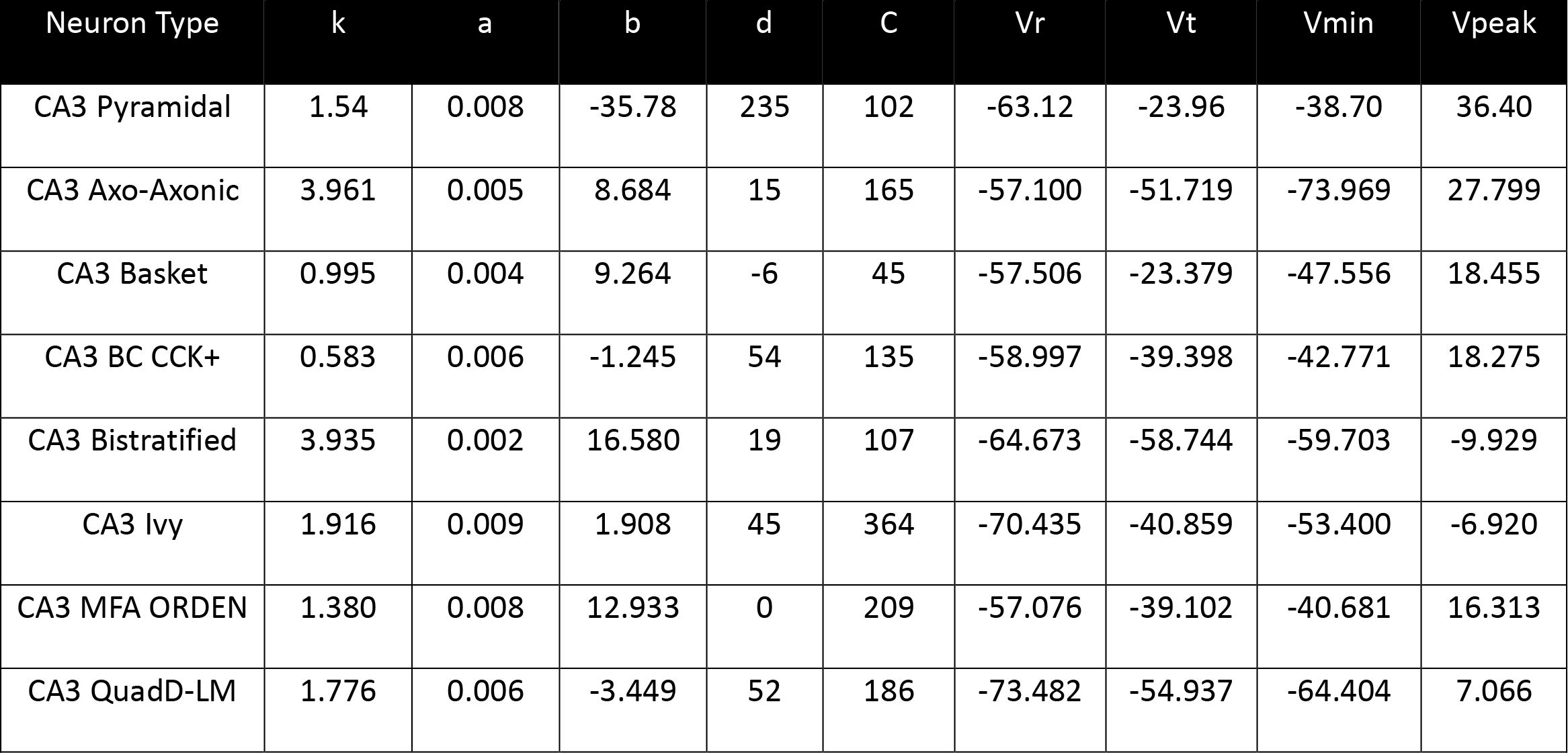
Izhikevich parameters by neuron type.

**Table 2.**
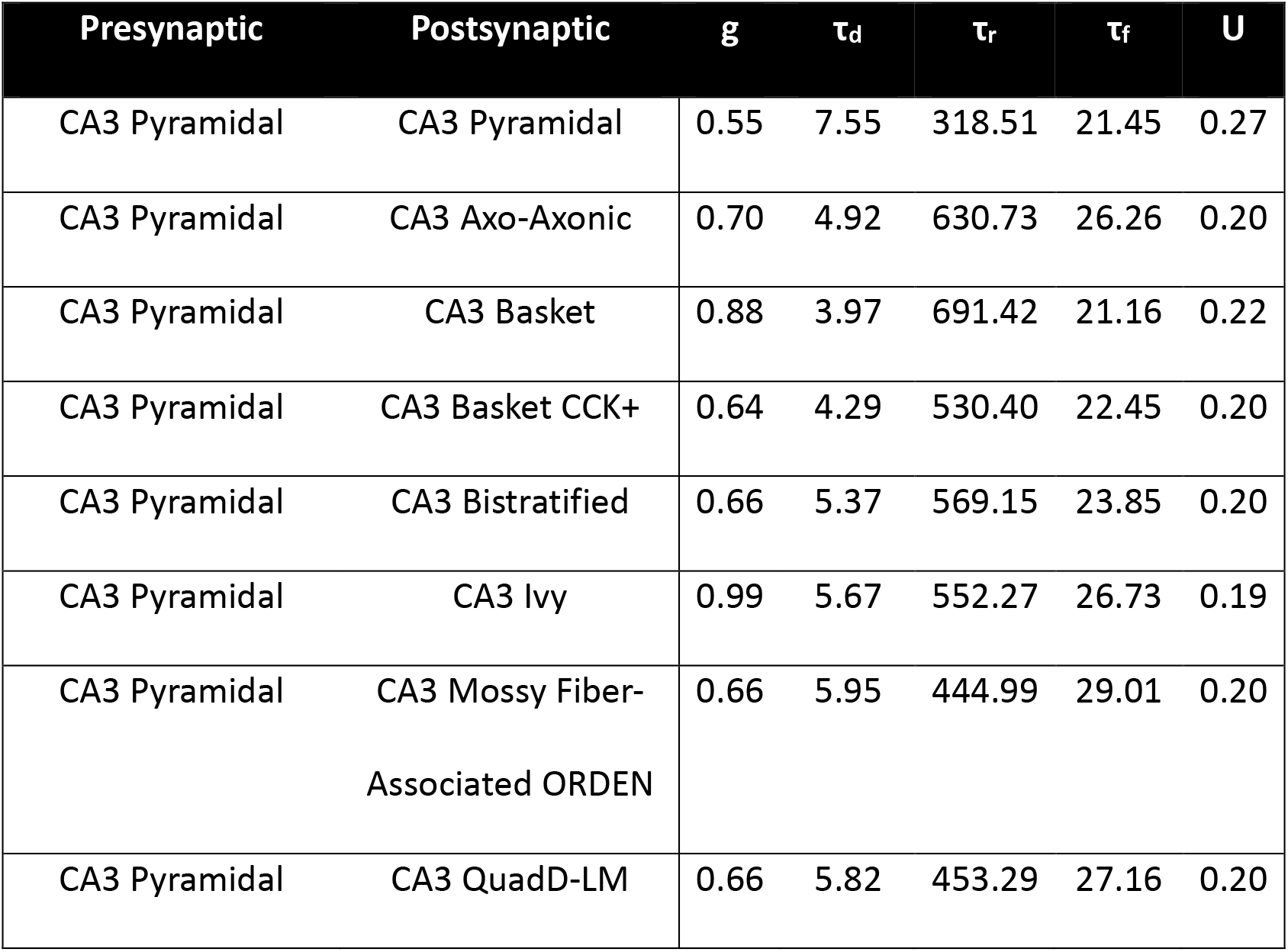
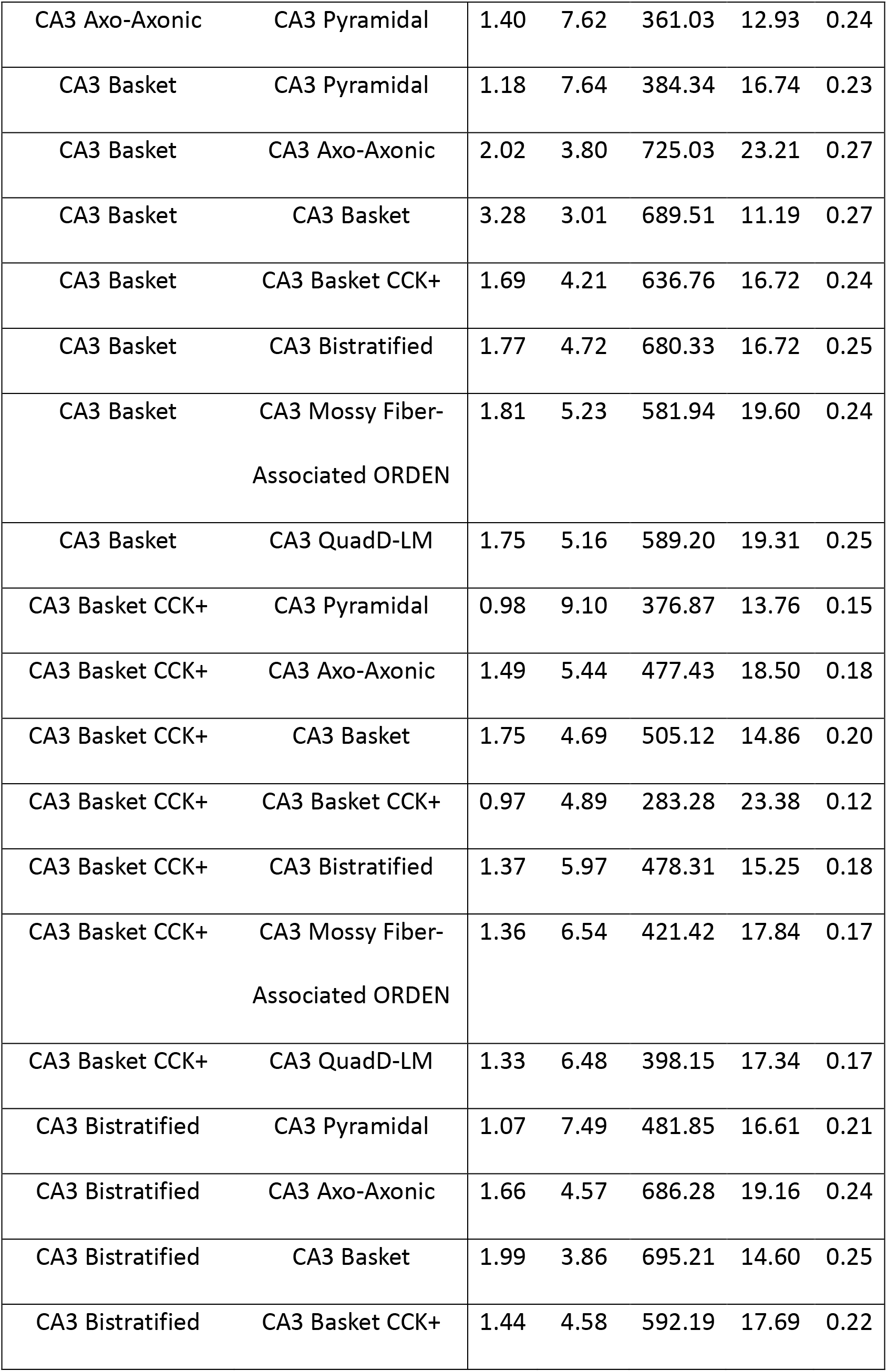
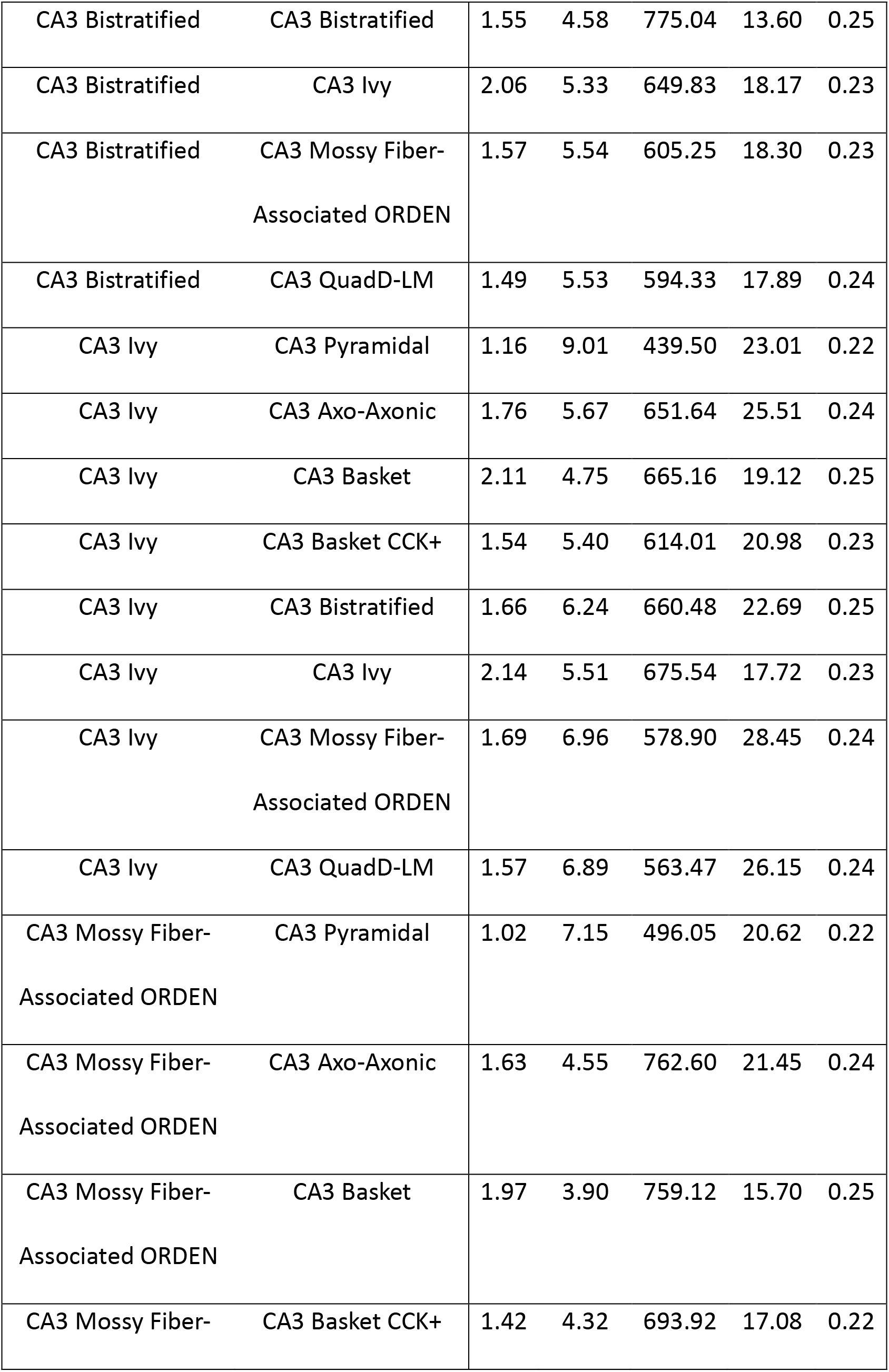
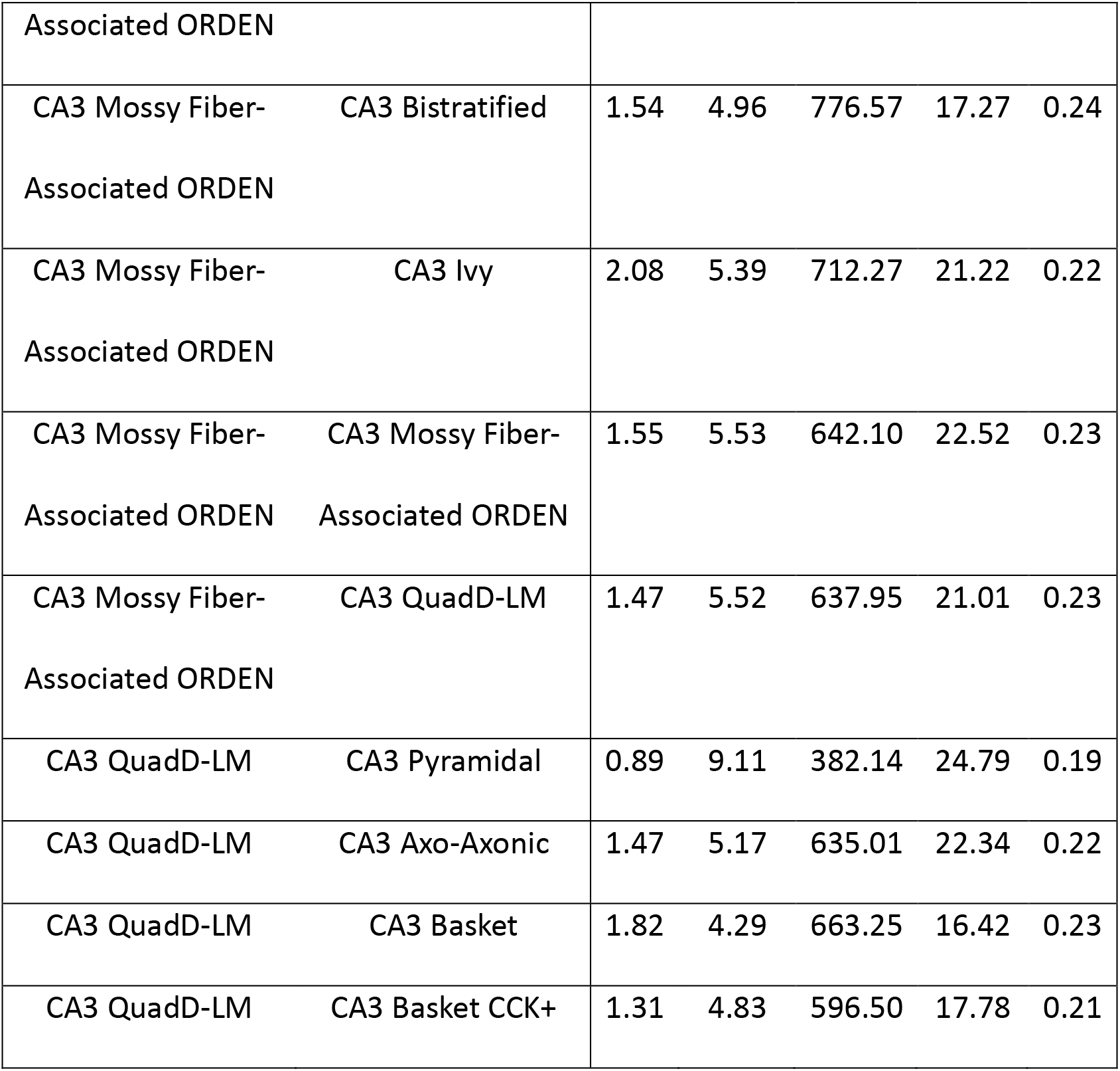
Tsodyks-Markram parameters for each connection type in the model.

We presented input patterns during a training phase that elicited concomitant firing in distinct subsets of PCs. This approach was inspired by a recent study [20] which demonstrated through functional connectivity analysis and network modeling that cell assemblies formed within CA3 from the application of different input patterns to subsets of CA3 PCs. In this work, each pattern lasted the length of a gamma cycle (20 ms) and was activated within an overarching theta cycle (200 ms), inspired by how cell assemblies are theorized to form *in vivo* according to a theta-gamma neural code ([7,40]; Figure 2a,c). After training, a degraded form of each input pattern was provided during a testing phase to evaluate the pattern completion capability of the SNN. Pattern degradation consisted of eliciting concomitant firing in a smaller subset of PCs than the subset used during training; the test consisted of ascertaining whether this subset could retrieve the full pattern during the second half of the theta cycle through activation of recurrent PC connections (Figure 2b,d).

**Figure 2:**
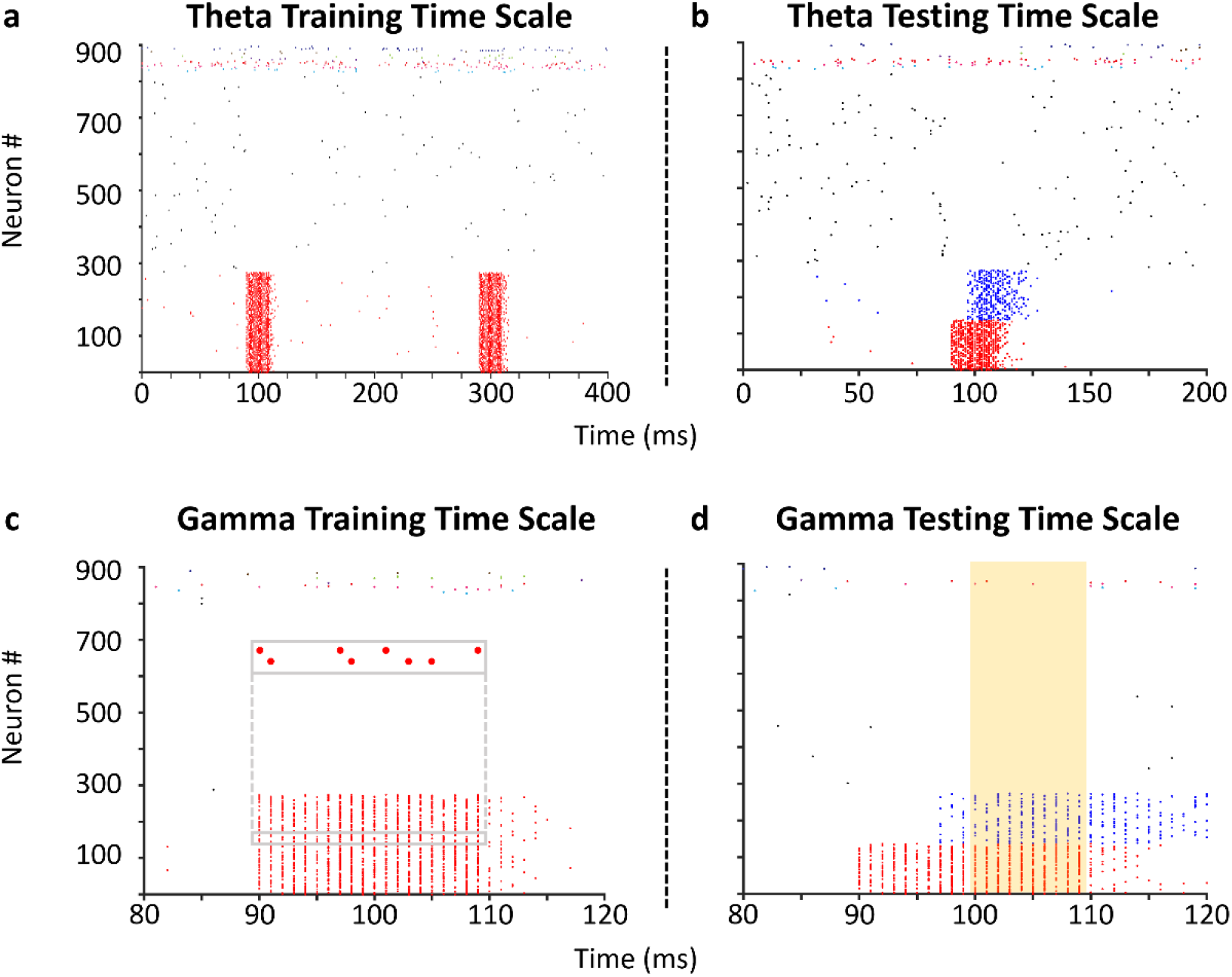
A theta-gamma training and testing protocol to investigate pattern completion within the CA3 SNN. (a) Training the SNN to store patterns involves the concomitant firing of (in this example) 275 PCs (red) during a theta time window. Two repetitions of a pattern are shown. Activity from a random selection of 500 PCs (black) and 10 interneurons of each type (spikes colored neither red nor black) are also shown. (b) Testing pattern completion involves only activating a subset of PCs during a theta time window. (c) Concomitant firing of PCs in (a) occurs during 20 ms gamma time windows. Inset: sparse firing of two representative neurons during pattern presentation. (d) Activation of a subset of PCs (red) in (b) leads to pattern completion of the remaining subset (blue) during a 20 ms gamma time window. The time window utilized for computing pattern reconstruction accuracy is highlighted by a gold rectangle (Supplementary Figure 2).

The full-scale network exhibited asynchronous population activity while patterns were not presented, with each neuron type firing at rates consistent with those observed for these types *in vivo* (Table 3). When patterns were presented, sparse firing of PCs was relegated primarily to assembly members, while the activity of each interneuron type remained similar to non-presentation periods (Figure 3a,c). Between training and testing, all PC-PC synaptic weights were re-normalized via synaptic divisive downscaling based on the synaptic homeostasis hypothesis [41–43]. In order to test the specificity of auto-association and pattern completion, we trained the network with three distinct input patterns. Training (with 65 repetitions in this example) induced strong auto-association through the synaptic weights of PCs within the subset of PCs stimulated by each input pattern, thereby forming three cell assemblies. Synaptic weights between members of different assemblies and between PCs that did not belong to any assembly were similar to the synaptic weights before training had commenced (Figure 3b). Strong auto-association within a subset of PCs stimulated by a given input pattern is indeed consistent with and expected from the cell assembly theory ([44]; Supplementary Figure 1). Stimulation of 50% of the input patterns provided during training (50% pattern degradation) led to robust activation of each assembly (Figure 3d).

**Figure 3:**
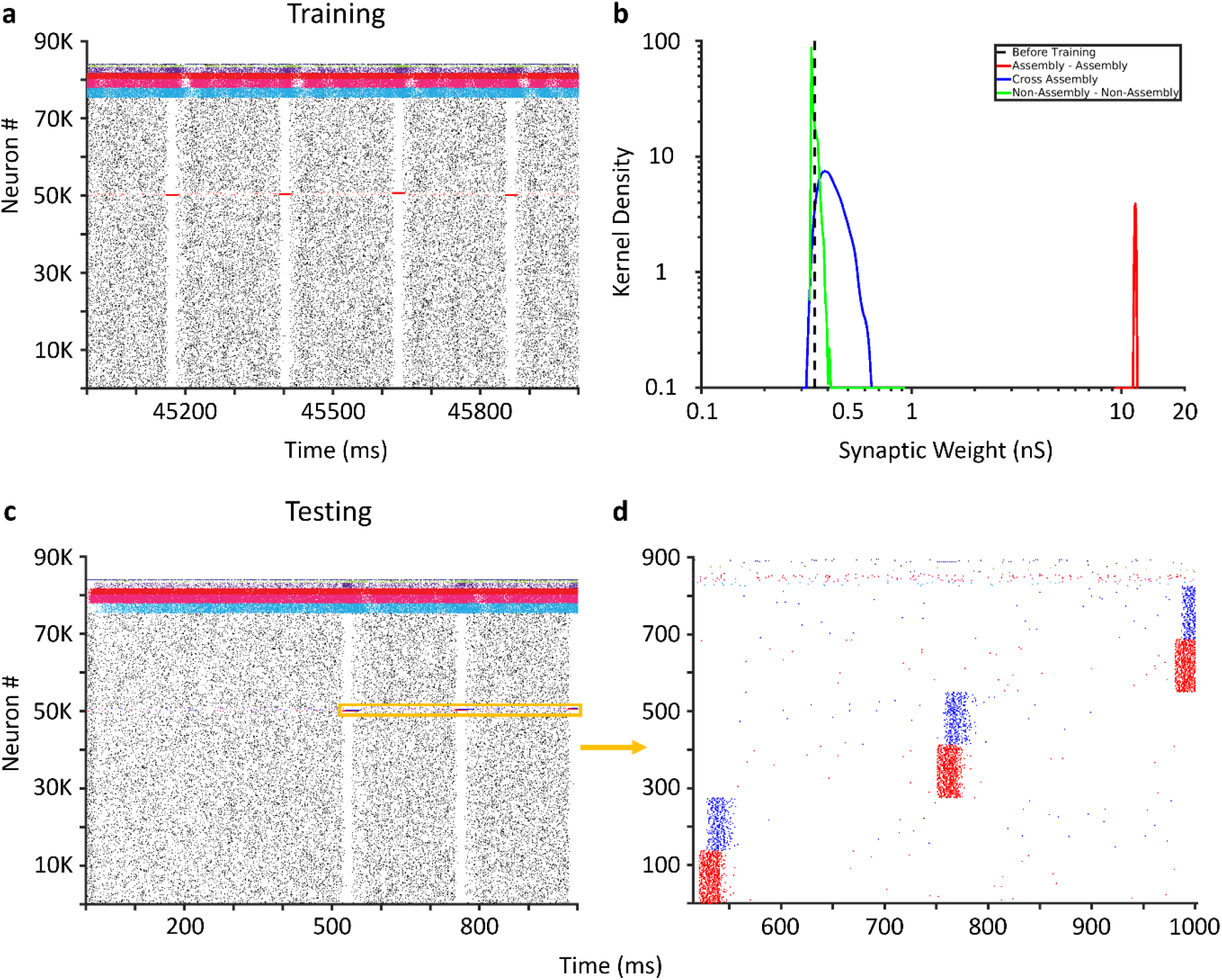
Pattern completion in the CA3 SNN. (a) Activity from the entire CA3 SNN during one second of training. (b) Kernel density estimates of PC-PC synaptic weights (after synaptic downscaling) within assembly (red), between members of different assemblies (blue), between non-assembly members (green), and the initial (uniform) synaptic weights before training (dashed black). (c) Activity from the entire CA3 SNN during one second of testing the recall of three patterns. Degraded patterns are presented at the five hundred millisecond mark (orange window). (d) Activity from 825 Pyramidal cells (PC) and 10 interneurons of each interneuron type (spikes that are neither red nor blue) during the orange window in (c). Input to 138 PCs (50% pattern degradation) in each assembly (red) leads to robust activation of the remaining assembly members (blue).

**Table 3.**
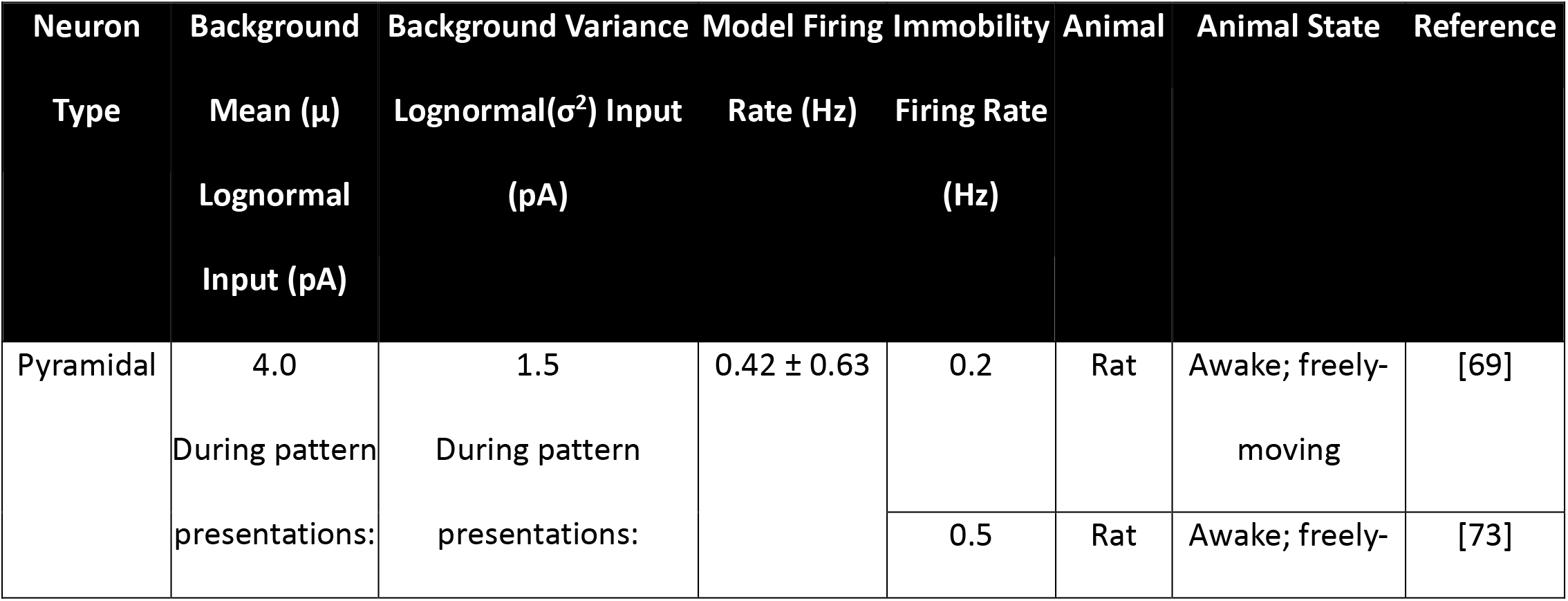
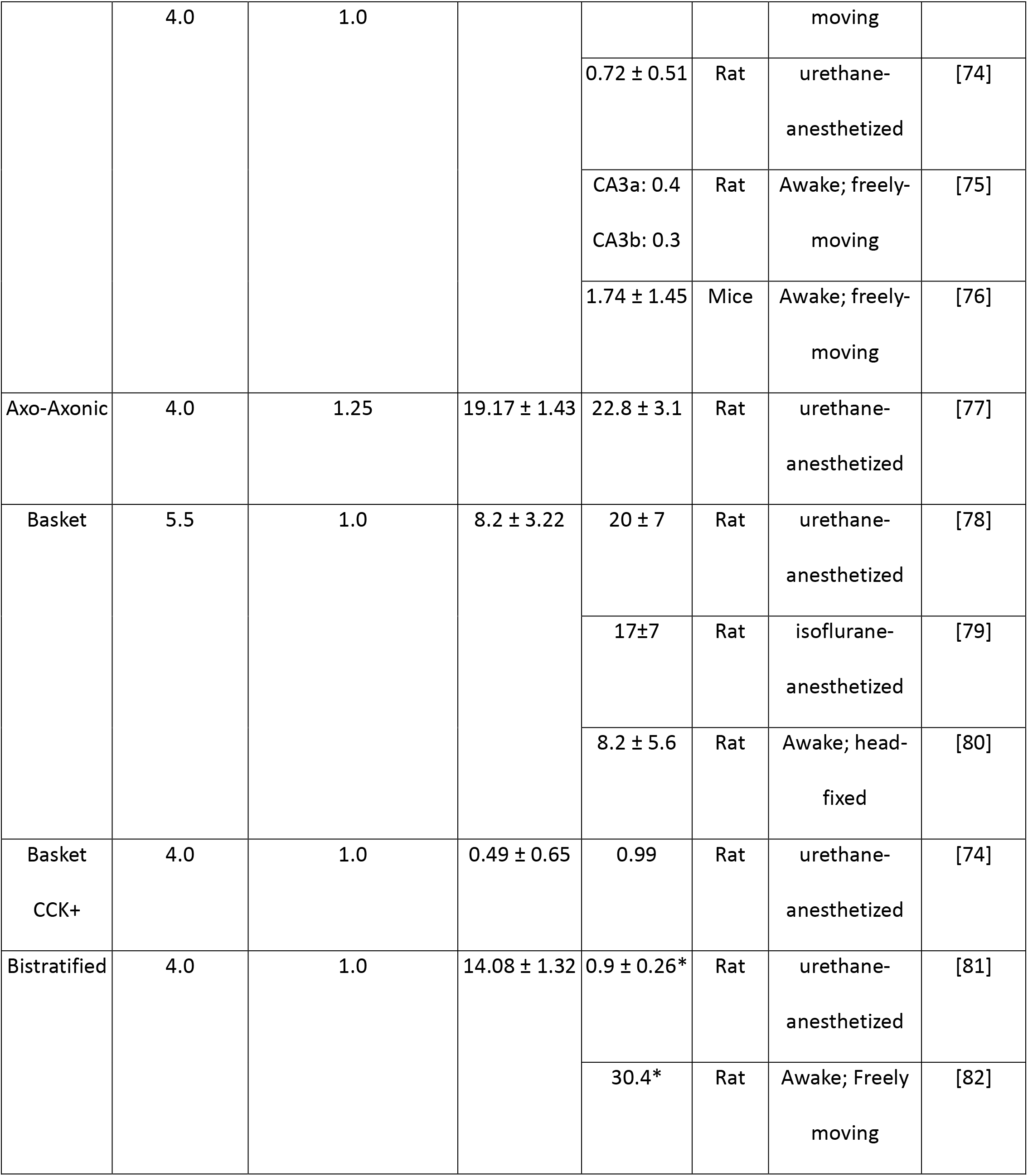
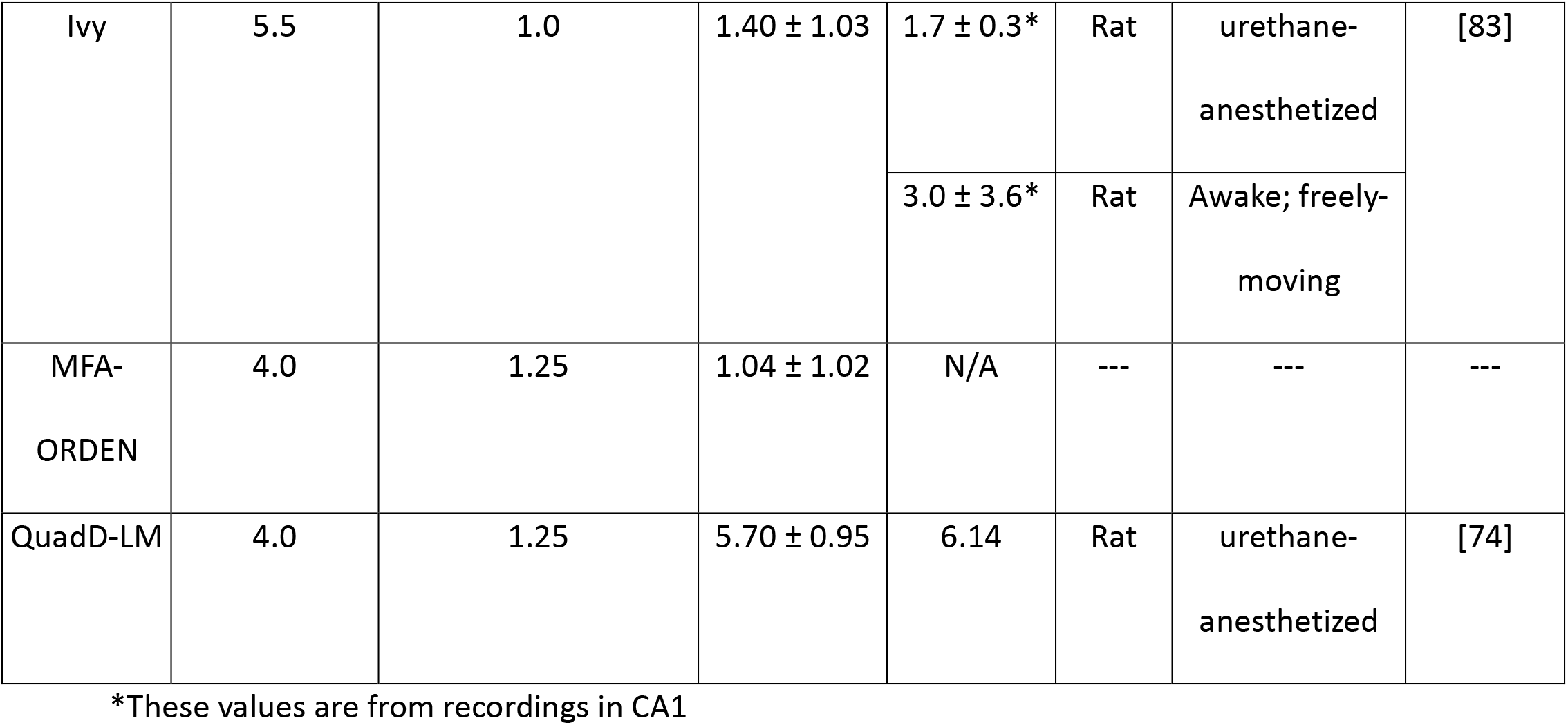
Firing rates (mean± s.d.) for each neuron type as recorded in our model and *in vivo*.

In summary, we extended a previous data-driven, full-scale SNN of the mouse CA3 with experimentally-derived STDP and showed that (1) the network could store patterns via cell assemblies when trained with a biologically realistic stimulation protocol; and (2) cell assemblies retrieved their activity patterns when only provided a halved cue. This result allowed us to investigate the robustness of cell assembly retrieval across a variety of scenarios.

### Can robust cell assembly retrieval occur across learning and with increasingly degraded cues?

A CA3 SNN capable of pattern storage and retrieval allows the characterization of two central aspects of auto-associative memory: the amount of repetition (learning) required for an experience to be stored and appropriately recalled, and the impact on performance when cues are degraded. Addressing these issues requires a metric to quantify the extent of pattern recall. To this aim, we defined *pattern reconstruction* based on a previously developed approach [45] relying on Pearson correlation coefficients (PCCs): if the output pattern PCCs were greater than the input pattern PCCs, then pattern completion occurred (Materials and Methods). Our pattern reconstruction metric adapts this index to capture the degree of pattern completion by scaling the PCCs relative to the maximum value of 1, and converting the result to a percentage to obtain an intuitive expression of performance accuracy (Supplementary Figure 2).

With pattern reconstruction defined, we turn to the first question. We trained the CA3 SNN in sets of 5 presentations of, again, three distinct input patterns, which in the prior example created three corresponding cell assemblies. After each set of 5 presentations, we stored the synaptic weight matrices of the network to enable separate testing with 50% degraded input patterns. Interestingly, non-zero pattern reconstruction occurred with as few as 15 presentations of input patterns (Figure 4a). Based on the second derivative of the reconstruction accuracy, 40 pattern presentations corresponded to the inflection point of most effective learning. Furthermore, a pattern completion plateau emerged at 55 presentations, with 65 and 95 presentations providing the strongest reconstruction accuracies, indicating the best pattern retrieval.

**Figure 4:**
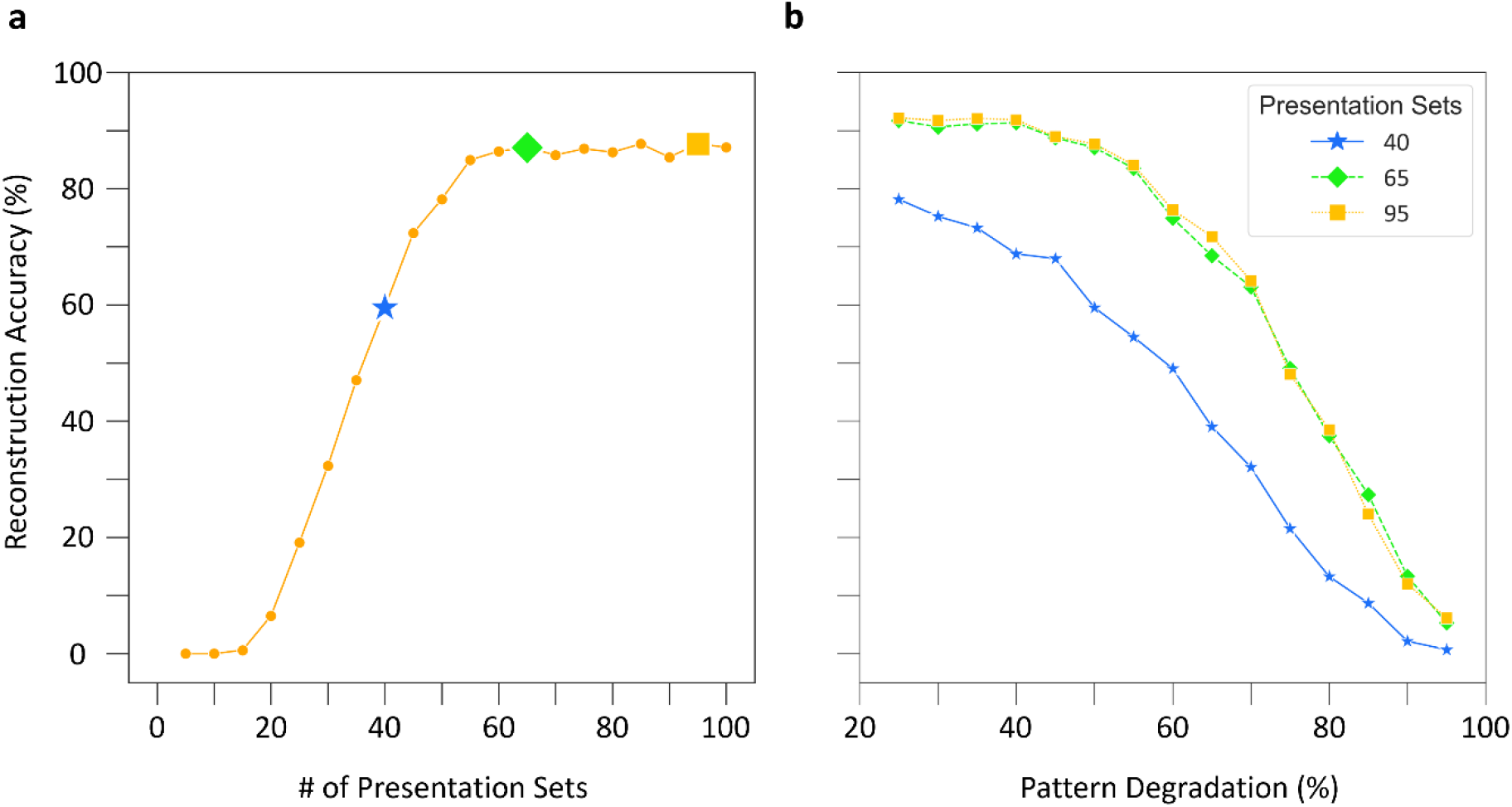
The CA3 SNN is robust to pattern degradation across learning. (a) Pattern completion accuracy, quantified by pattern reconstruction with 50% pattern degradation, as a function of training. The star denotes the inflection point for most effective learning as defined by the second derivative of the accuracy curve, and the diamond and square denote the two best accuracy values on the plateau. (b) Reconstruction accuracy as a function of pattern degradation. With increased training, cell assemblies can withstand greater degradation of input patterns, but only up to the initial plateau. Results in both panels are from an assembly size of 275.

Turning to the second question, we utilized network structures trained on 40, 65, and 95 pattern presentation sets to assess how increased pattern degradation (i.e., increasingly diminished pattern cues) impacted pattern retrieval. Remarkably, pattern reconstruction remained substantial until a steep drop-off at 70% pattern degradation, and only weak pattern reconstruction occurred with 95% pattern degradation for each of the three network structures (Figure 4b). Additionally, the similar performance of networks trained on 65 and 95 input repetitions highlighted that training beyond the initial plateau does not improve performance at more extreme pattern degradations.

Taken together, these results show that the CA3 SNN reliably encoded and retrieved patterns after as few as 40 presentation sets and upon reactivation of only a minority of PCs belonging to a cell assembly.

### What assembly sizes can support pattern completion?

Another fundamental question is that of memory capacity – how many experiences can the network store and recall without interference? To address this question, we first consider the simple scenario in which all cell assemblies are fully segregated, that is, no neuron belongs to more than one assembly. In this case, the number of cell assemblies supported by the CA3 network is given by the total number of CA3 pyramidal cells divided by the assembly size, i.e. the number of CA3 pyramidal cells constituting each assembly. This factor is related to the sparseness ratio (γ), defined as the percentage of cells activated during an experience [19]. Theoretical insights and experimental evidence from humans and rats offered constraints for γ; using these constraints as a guide, we tested cell assembly sizes between 50 and 600 (0.067% <= γ <= 0.8%; [19,20,46,47]; Materials and Methods).

We trained networks on 40, 65, and 95 presentation sets to create assemblies of variable size and tested on patterns degraded by 50%. Interestingly, smaller sized assemblies performed best with fewer presentations (40 sets), while larger sized assemblies performed best with more presentations (65 and 95 sets) (Figure 5a). Additionally, there was a stable range of assembly sizes between 150 and 600 where reconstruction accuracy improved with more training; the best performance occurred for an assembly size of 275. Assembly sizes smaller than 150 with additional training performed worse due to pattern interference (Supplementary Figure 3). Furthermore, as observed in the previous section, the choice of either 65 or 95 presentation sets within this range conferred similar pattern reconstruction accuracy.

**Figure 5:**
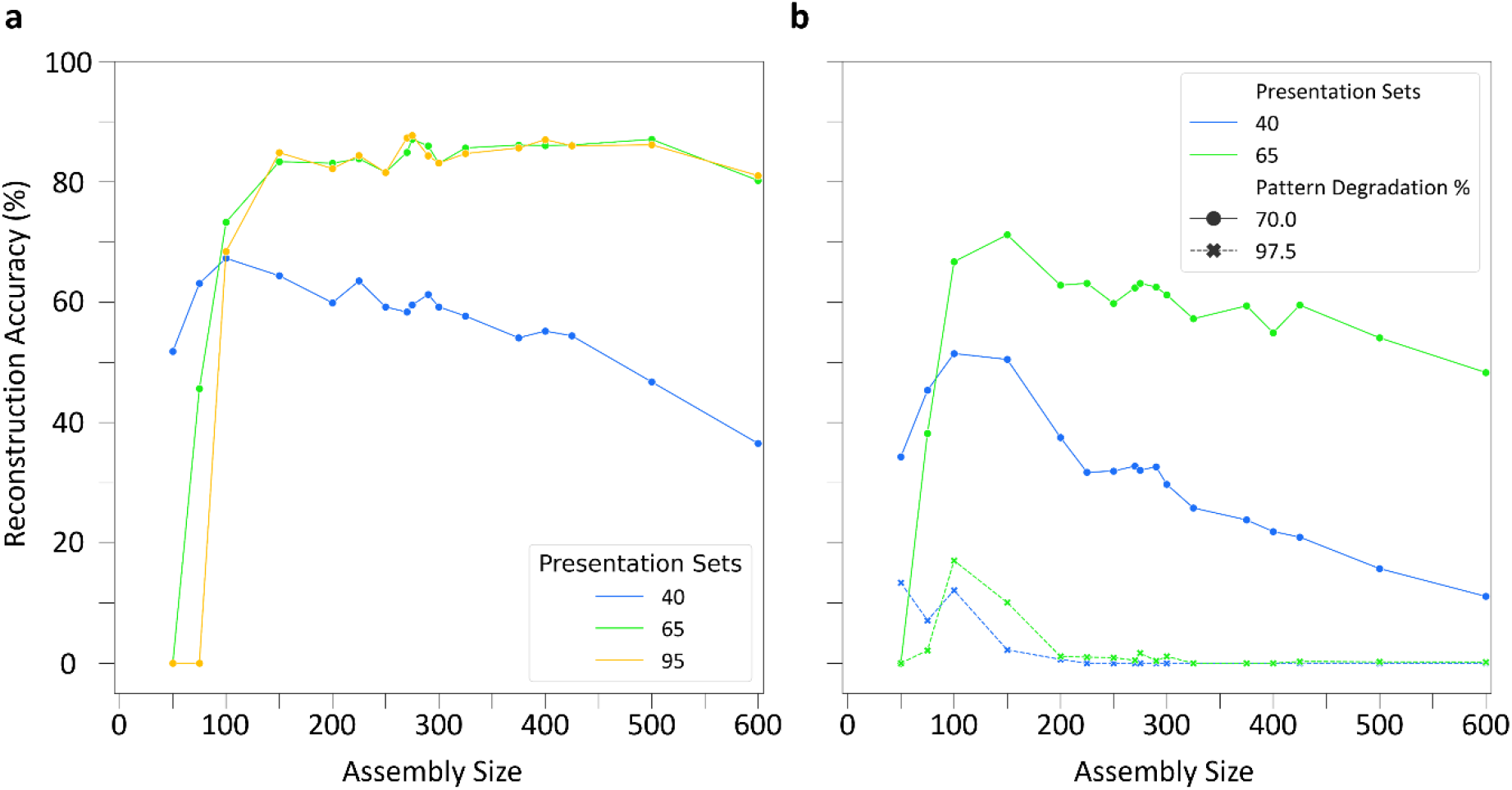
A range of cell assembly sizes can support robust pattern completion. (a) Reconstruction accuracies for a range of cell assembly sizes throughout learning with 50% pattern degradation. (b) Effect of assembly size on reconstruction accuracy with increased pattern degradation levels of 70 and 97.5%.

Utilizing SNNs trained on 40 and 65 presentation sets, we further tested pattern completion for the range of assembly sizes with increased pattern degradation percentages of 70 and 97.5% (Figure 5b). Notably, assembly sizes of 100 and 150 displayed the best pattern completion in response to these highly degraded input patterns and exhibited weak pattern completion even when only 2.5% of an input pattern was provided. Therefore, in the presence of severely degraded input patterns, smaller assembly sizes (100 and 150) performed best in the SNN, whereas across moderate to high degradation levels an assembly size of 275 offered the best performance.

### Can a full-scale CA3 SNN store and recall overlapping cell assemblies?

Our analysis so far assumed that no neuron could belong to more than a single cell assembly, but this is not necessarily the case in biological circuits. In fact, the extent of assembly overlap constitutes another key factor in determining memory capacity, because sharing neurons between cell assemblies can increase the number the experiences the network can encode [22]. Moreover, neurons shared between cell assemblies may facilitate hetero-association between episodic memories in CA3 [21]. Therefore, we investigated the storage and retrieval of patterns in the CA3 SNN when cell assemblies shared a subset of neurons (Figure 6a).

**Figure 6:**
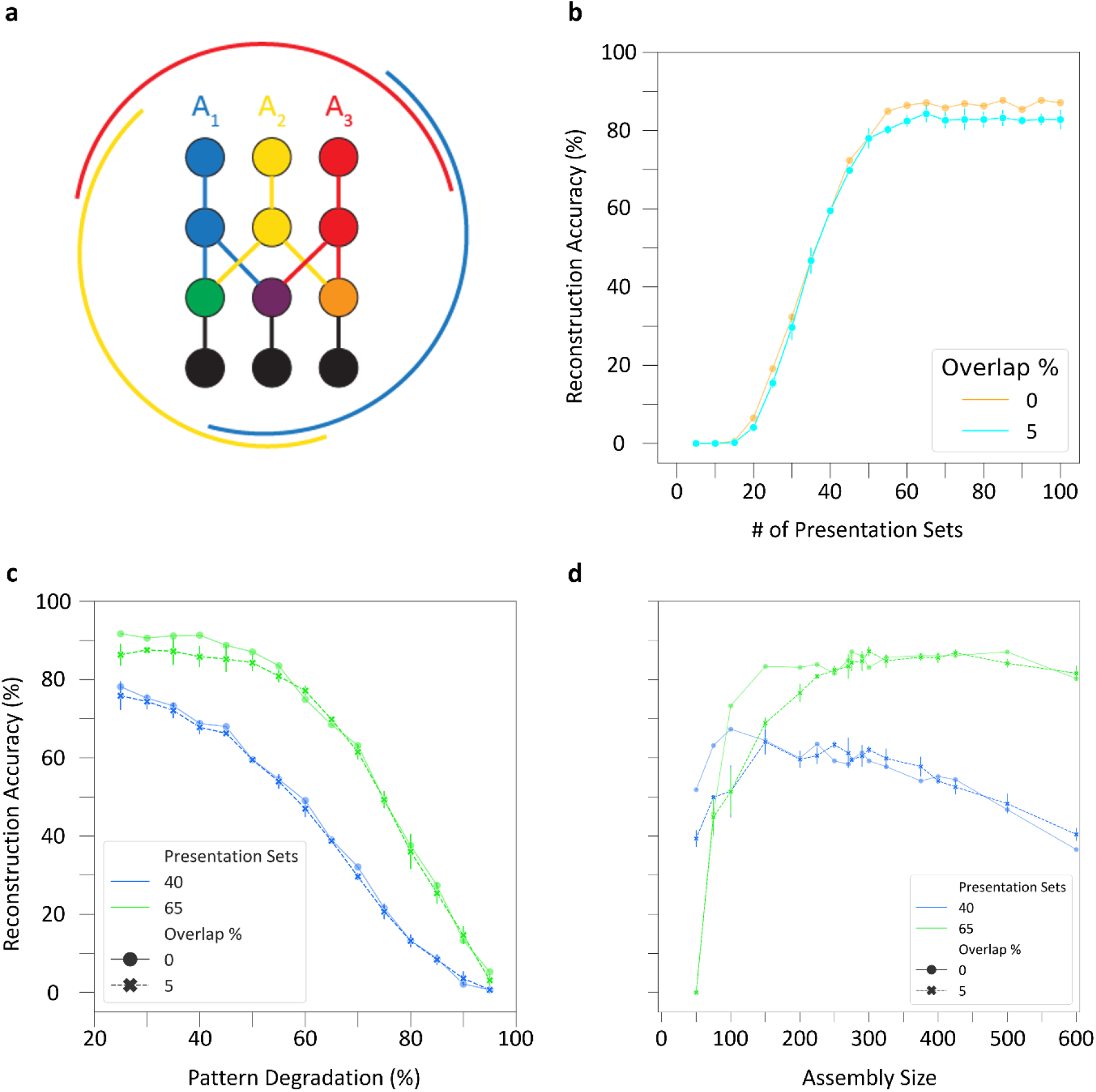
Overlapping cell assemblies support robust pattern completion. (a) Schematic of cell assembly overlaps. Three assemblies (red, blue, and yellow) of neurons (circles) and connections (lines), with shared cells (green, purple, and orange circles). Black circles and lines represent non-assembly neurons. The external arcs indicate the extent of overlaps. (b) Overlapping cell assemblies display similar reconstruction accuracy to assemblies without overlap throughout learning at 50% pattern degradation. (c) Overlapping cell assemblies perform comparably in reconstruction accuracy to assemblies without overlap when pattern degradation is increased. Results from (b) and (c) are from an assembly size of 275. (d) Overlapping cell assemblies have comparable reconstruction accuracy to assemblies without overlap across a range of assembly sizes at 50% pattern degradation. Bars in (b-d) reflect standard deviation of accuracy across three simulations with randomized selection of overlapping cells.

To create overlaps between the three cell assemblies, we randomly selected 5% of neurons as shared between each pair of assemblies before training commenced (Materials and Methods). Following the usual procedure for storing cell assemblies and degrading input patterns by 50% during testing, overlapping cell assemblies retrieved patterns comparably to cell assemblies without overlaps (Figure 6b). In particular, pattern reconstruction accuracy followed a similar trajectory with overlapping cell assemblies and had the same optimal point for learning of patterns and highest accuracy, which occurred at 40 and 65 presentations, respectively. Additionally, testing the overlapping cell assemblies in the presence of increased pattern degradation after training with 40 and 65 pattern presentation sets yielded similar reconstruction accuracies as with the no overlap (Figure 6c). Furthermore, in the presence of 5% overlap, cell assembly sizes between 200 and 600 supported strong pattern completion, again consistent with the range found for assemblies without shared cells (Figure 6d); notably, however, overlap reduced the performance of smaller cell assemblies in the 50-150 range.

Auto-association and pattern completion of cell assemblies reflect the structural and functional components of memory formation and recall within the CA3 circuit, respectively, and SNNs can help reveal the underlying link between structure and function [7,48]. We investigated this relationship by tracking two characteristics of PC-PC synapses throughout training: the auto-association signal-to-noise ratio (SNR) and the percentage of assembly synapses that had reached the maximum weight (Materials and Methods). It is especially interesting to analyze if and how these characteristics relate to the observed pattern completion performance. In this regard, we observed an auto-association SNR plateau occurring in assemblies trained both with and without overlap: further improvements in reconstruction accuracy became inconsequential above 94% SNR (Figure 7a). This is consistent with the influence of the number of presentations on pattern completion, where training beyond 60 presentation sets did not significantly improve retrieval (cf. Figure 6b). Furthermore, at both 50% and 70% pattern degradation, reconstruction accuracy reached values close to optimal performance when only 10% of assembly synapses had reached their maximum weight with or without overlap (Figure 7b). This indicates that effective learning in the CA3 SNN does not require synaptic saturation.

**Figure 7:**
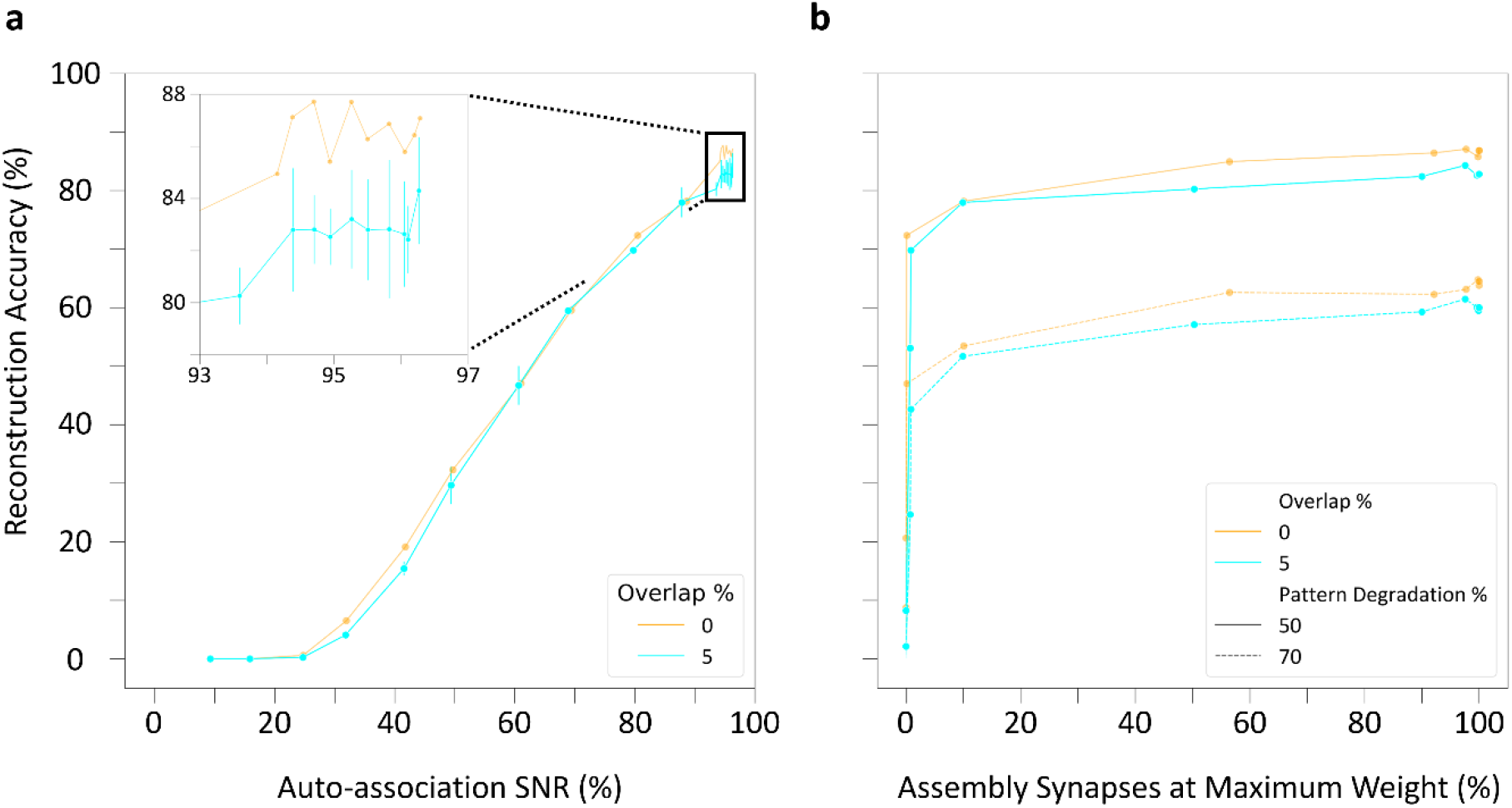
Relationship between pattern completion performance and synaptic characteristics of the CA3 SNN. (a) Auto-association of cell assemblies throughout learning highlights comparable reconstruction accuracy as a function of maximum signal-to-noise ratio (SNR) regardless of overlap percentage. Inset: Zoomed-in view near maximum auto-association SNR. (b) Pattern completion as a function of the percentage of synapses at maximum weight demonstrates that optimal performance does not require synaptic saturation.

Taken together, these results highlight that strong retrieval occurs at moderate SNRs and when most assembly synapses are below their maximum weight. Moreover, overlapping cell assemblies retrieve patterns comparably to non-overlapping assemblies, supporting the use of shared neurons to enhance auto-associative memory capacity.

## Discussion

The present work demonstrates that a biologically realistic SNN of the mouse CA3, with cell type-specific parameters of neuronal excitability, connection probabilities, and synaptic signaling all extracted from experimental measurements, can store and recall auto-associative memories via cell assemblies. Notably, cell assembly formation and retrieval relies on a training and testing paradigm grounded by *in vivo* neurophysiology [40]. In particular, strong pattern reconstruction reliably occurs in the SNN in response to heavily incomplete or degraded input cues. Furthermore, auto-associative pattern completion in our model is robust across a broad range of assembly sizes and in the presence of assembly overlap, two critical factors to determine auto-associative memory capacity in CA3.

Training our CA3 SNN to an optimum point enabling strong pattern completion, yet well before most assembly synapses reach their maximum weights, may reflect how the real CA3 stores and recalls memories. Rather than maximizing post-synaptic conductance in pyramidal cells, a tradeoff with synaptic downscaling (possibly during slow-wave sleep) could support the storage of many patterns with minimal pattern interference. Additionally, these insights may be useful in training artificial neural networks, where training a network on multiple tasks in parallel with reasonable performance, instead of optimizing accuracy on a single task, could prevent catastrophic forgetting [49,50].

The hippocampus may facilitate “one-shot” learning, i.e. rapid memory encoding from just a single experience [51]. In a previous study, training a rat CA3 network model to store patterns with a clipped Hebbian plasticity rule enabled encoding of these memories in a one-shot manner [20]. However, one-shot encoding may not be prominent in the real rodent hippocampus, as animals typically spend weeks to learn a spatial or novel object location memory task before testing begins, and even then strong performance often requires multiple trials [2,52–55]. During this time the hippocampus goes through many encoding, consolidation, and retrieval phases, when theta, gamma, and sharp-wave ripples contribute to cell assembly formation, refinement, and recall [56]. Therefore, our simulation design subjected the CA3 SNN to a training phase representative of encoding during experience through theta nested gamma oscillations [40]. Moreover, modification of synaptic weights within assemblies between training and testing reflected synaptic downscaling during slow-wave sleep [57]. With this protocol, heavily degraded or incomplete cues reliably triggered strong pattern completion-mediated recall of experiences, in line with the expected role of CA3 in auto-associative memory.

Our results of robust pattern completion using circuit parameters measured from anatomical and physiological experiments complement and extend previous modeling work. A network model consisting of CA3 PCs and two interneuron types receiving inputs from the entorhinal cortex and dentate gyrus showed that, when patterns were strongly degraded, pattern completion could still occur within one recall cycle, known as simple recall [47,58]. Neither this model nor the rat CA3 network model mentioned above [20], however, constrained the simulation based on both size and diversity of the CA3 circuit. Another recent model of pattern completion in CA3 reflected the mouse network size, but again not the neuronal and synaptic diversity [59]. Therefore, to our knowledge this work provides clear evidence of robust pattern completion in the most realistic SNN model of the mouse CA3 to date.

The cell assemblies formed and retrieved in this work involved either zero or 5% shared cells between them. It is likely that cell assemblies have at least *some* level of overlap between them, as randomly creating assemblies with coding sparseness ratio of γ would share γ^2^N cells in common [21]. Our results with substantial overlap demonstrate that the neuronal and synaptic physiology of the CA3 circuit are well suited to support pattern completion even if non-zero overlap exists between assemblies in the mouse hippocampus *in vivo*, as recent empirical evidence demonstrates in the mouse primary visual cortex [60].

The best performance across the range of assembly sizes examined in this study, when considering varying levels of cue degradation, lengths of training, and overlap, occurred with an assembly size of between 250 and 300 neurons. Intriguingly, 275 is approximately the square root of the number of PCs in the mouse CA3 network. It is tempting to speculate that hippocampal cell assemblies *in vivo* optimally form in accordance with the square root of the number of PCs, at least in rodents: based on the values of γ reported in previous studies, the square root relation would hold for rats, but not for humans. However, these estimates for γ are based on indirect evidence, including the number of hippocampal place cells active in each environment, and the number of concept cells active when presenting a concept, based on simultaneous single- and few-neuron recordings [19,61].

Estimation of memory capacity in CA3 has previously involved the use of the connection probability, c, between CA3 PCs and γ. Utilizing Willshaw’s formula, which estimates capacity for both non-overlapping and overlapping assemblies with P = c/γ^2^, the capacity of the mouse CA3 would be on the order of 2,000 patterns [19]. Another formula was proposed by Treves and Rolls, which considers the number of recurrent collateral (RC) connections onto each PC, C^RC^, a scaling factor reflecting the total amount of information that can be stored and retrieved from the RCs, k, and γ [5,23]. Estimating capacity with their formula of 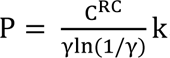, the mouse CA3 could store on the order of 18,000 patterns. However, these formulas do not consider overlap directly as a variable, which may in principle allow a substantial increase in storage capacity.

The advent of large-scale recording technologies, including two-photon calcium imaging, Neuropixel probes, and hundred Stimulation Targets Across Regions (HectoSTAR), enabling the simultaneous monitoring of thousands of neurons, may soon make it feasible to measure more directly the size of hippocampal assemblies [62–64]. Such evidence might show that the size of assemblies *in vivo* could vary depending on the represented cognitive content, providing further guidance for how to extend our SNN model. Additionally, these recordings during cue mismatch tasks would pinpoint how many neurons are typically reactivated in response to degraded cues [14,65], allowing a quantitative comparison with our results. Last but not least, large-scale recordings will likely highlight the variation in neuronal overlap between assemblies, facilitating the estimation of key factors determining the memory capacity of the CA3 circuit.

## Materials and Methods

### Full-scale CA3 SNN

The selection of the neuron types constituting the CA3 SNN and the model parameters, including neuron type-specific excitability, population size, connection probabilities, and synaptic signaling, were developed and validated in prior work [38]. Briefly, the SNN consists of PCs and seven interneuron types: Axo-axonic, Basket, Basket CCK+, Bistratified, Ivy, Mossy Fiber-Associated ORDEN (MFA-ORDEN), and QuadD-LM cells. The perisomatic targeting and axonal-dendritic overlaps between these eight neuron types give rise to 51 directional connections (Figure 1a).

For each neuron type, we utilized experimentally-derived parameters from Hippocampome.org for both the neuronal input-output function, i.e., the spiking pattern produced in response to a given stimulation, and the neuron count. In particular, to balance biological realism with computational efficiency, we chose the Izhikhevich 9-parameter, single-compartment dynamical systems framework [66]. The parameters reflect the following neuron type-specific properties: membrane capacitance (*C*), a constant that reflects conductance during spike generation (*k*), resting membrane potential (*vr*), instantaneous threshold potential (*vt*), a recovery time constant (*a*), a constant that reflects conductance during repolarization (*b*), spike cutoff value (*vpeak*), reset membrane potential (*Vmin*), and a constant that reflects the currents activated during a spike (*d*). Hippocampome.org reports the parameter values that best fit the firing patterns reported in the literature for the corresponding neuron types [36].

For neuron counts, we considered each neuron type in our network as a representative of its *supertype* family (hippocampome.org/morphology). Thus, the population size of each neuron type in the SNN is the sum of all neuron types of the given supertype. For example, the number of instantiated CA3 Axo-axonic cells in the model (i.e., the population size parameter value for this particular neuron type) consisted of the sum of Axo-axonic proper and Horizontal Axo-axonic cells (two variants of Axo-axonic neurons in CA3), which Hippocampome.org reports as 1,482 for the mouse. The population sizes and the 9 Izhikevich parameters for each of the 8 CA3 neuron types are shown and listed in Figure 1a and Table 1, respectively.

Modeling neuron type-specific communication involves a description of the postsynaptic signal caused by a presynaptic spike and related short-term plasticity (STP), as well as the connection probability and delay between the presynaptic and the postsynaptic neuron types. We modeled synaptic dynamics with the 5-parameter Tsodyks-Markram framework [67], for which Hippocampome.org reports experimentally-derived pre- and post-synaptic neuron type-specific values (Table 2): synaptic conductance (g), decay time constant (τd), resource recovery time constant (τr), resource utilization reduction time constant (τf), and portion of available resources utilized on each synaptic event (U). Note that this formalism captures *unitary* synaptic communication. As such, it reflects the total somatic effect of all synapses corresponding to connected neuron pairs. Given the local scope of the CA3 circuit, all connections were modeled with a synaptic delay of 1 ms. Hippocampome.org also provides morphologically derived connection probabilities for each directional pair of rat neuron types [35], which we scaled for the mouse according to a fixed anatomical sizing ratio [68]. The probabilities for all 51 connection types in the circuit are reported in Figure 1a.

Every instantiation of the simulation thus contained 84,053 neurons and 176 million synaptic connections on average. To elicit activity in the SNN, each neuron received a lognormal background current to model the upstream inputs CA3 receives from dentate gyrus and entorhinal cortex [69,70]. The inputs were constrained to match the mean firing rates of each neuron type in the model with those observed *in vivo* (Table 3).

### Range of assembly sizes

In order to define a range of assembly sizes to evaluate auto-association and pattern completion, we first considered the sparseness ratio of neural coding, γ, which is the average fraction of cells activated during an experience [19]. Available estimates for γ in humans and rats varied only slightly, from 0.1% [20], through 0.23% [46], to 0.3% [19]. This would correspond, for the number of PCs in mouse CA3, to a range of sizes between 75 and 225. The authors of the latter cited study, however, accompanied their estimate for assembly size (225) with a wider range (150-300) as well as cautionary lower and upper bounds of a factor of 2 in either direction [14]. Furthermore, in the absence of precise experimental determinations, smaller values of value of γ could allow for larger storage capacity as long as recall from partial input could be maintained [47]. Based on these lines of reasoning, we set bounds of 0.067% <= γ <= 0.8%, corresponding to a range of assembly sizes between 50 and 600.

### Long-term synaptic plasticity

In line with the notion that cell assemblies form via long-term plasticity [71], we adopted a symmetric (Hebbian) spike-timing dependent plasticity (STDP) learning rule between PCs [12]: Δ*w* = *Ae*^−|Δ*t*|/*τ*^. Here, Δ*w* is the change in synaptic weight, *A* determines the weight change where the pre- and post-synaptic neurons fire at exactly the same time, *τ* is the plasticity decay time constant, and Δt is the temporal difference between the post- and pre-synaptic spikes. The value for *τ* was set to 20 ms, which best approximated the symmetric exponential decay curve observed experimentally for CA3 PCs [12] (Figure 1B). The values for *A* varied based on the maximum CARLsim6 synaptic weight (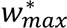) between PCs, which in our model depended on cell assembly size. Specifically, since the firing of each PC is triggered by the convergent integration of all activated presynaptic PCs, we reasoned that the maximum synaptic weight of each synapse should be inversely proportional to the number of PCs in an assembly.

In initial pilot testing with an assembly size of 300, we found that a value 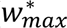 = 20 induced strong auto-association after 100 input pattern presentations. Therefore, we anchored the maximum synaptic weight scaling based on assembly size to this value: for instance, SNNs with assembly size of 150 or 600 would have a 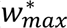 of 40 or 10, respectively. We then derived *A* so as to allow the synaptic weight to increase from the initial value before training (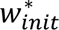 = 0.625 in all our simulations) to 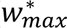 if all pre- and post-synaptic spikes were exactly coincident during training in the initial pilot settings. Since each of the 100 randomized spike trains during training contain 4 spikes on average, the resulting formula was 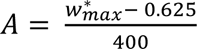, where 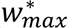. Table 4 reports the maximum total synaptic conductance (*g_max_* = 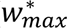 * *g*) and 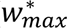 between PCs and *A* for each assembly size used.

**Table 4.** Maximum synaptic conductances, weights, and learning rates for each assembly size.

| Assembly Size | $g_{max}$ | $w_{max}^*$ | $A$ |
| --- | --- | --- | --- |
| 50 | 66.37 | 120 | 0.298 |
| 75 | 44.25 | 80 | 0.198 |
| 100 | 33.19 | 60 | 0.148 |
| 150 | 22.12 | 40 | 0.098 |
| 200 | 16.59 | 30 | 0.073 |
| 225 | 14.75 | 26.67 | 0.065 |
| 250 | 13.27 | 24 | 0.058 |
| 270 | 12.29 | 22.22 | 0.054 |
| 275 | 12.07 | 21.82 | 0.053 |
| 290 | 11.44 | 20.69 | 0.050 |
| 300 | 11.06 | 20 | 0.048 |
| 325 | 10.21 | 18.46 | 0.045 |
| 375 | 8.85 | 16 | 0.038 |
| 400 | 8.30 | 15 | 0.036 |
| 425 | 7.81 | 14.12 | 0.034 |
| 500 | 6.64 | 12 | 0.028 |
| 600 | 5.53 | 10 | 0.023 |

### Network training and testing protocol

Formation and retrieval of cell assemblies occurred in the CA3 SNN through dedicated training and testing phases. During the training phase, the SNN was presented with three input patterns, which consisted of requisite injected current to activate firing in a specific subset of PCs based on the size set for an assembly. The current injections triggered in each PC a randomized train of four spikes during a 20 ms (gamma) time window, with 200 ms (theta) time windows separating the presentation of the subsequent input pattern. This protocol of patterns presented at 50 Hz within an encompassing 5 Hz rhythm (“theta-gamma neural code” [7,40]) resulted in the formation of three unique cell assemblies. After the initial randomization of spike trains in the first input pattern, the same pattern was provided to each subset of PCs in every subsequent presentation of the pattern.

Between training and testing, each synaptic weight (*w*\*) between PCs was divided by the same factor such that the average *w*\* across all PC-PC synapses returned to 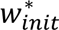. Rescaling synaptic weights in this manner is theorized to occur during slow-wave sleep, preserving synaptic weight distributions without eliminating the auto-association between assembly member PCs [41–43].

Testing pattern completion involved providing degraded input patterns to the SNN during gamma and theta time windows as performed during training. Degradation of input patterns consisted of decreasing the percentage of assembly PCs firing together within the designated 20 ms period. The percentage of pattern degradation in this work ranged from 25% to 97.5%.

### Quantification of auto-association and pattern completion

The capability of the SNN to form cell assemblies was investigated by quantifying two features of PC-PC synapses. The first one was the auto-association signal-to-noise ratio (SNR), defined as the mean synaptic weight between assembly member PCs divided by the mean synaptic weight between non-assembly member PCs. Thus, the higher the ratio, the stronger the auto-association of the formed cell assemblies relative to the rest of the CA3 network. The maximum auto-association SNR for each network structure investigated would occur if all assembly and non-assembly members had reached the downscaled maximum and minimum synaptic weights, respectively. The second quantified feature was the percentage of all assembly member synapses that had reached the maximum synaptic weight.

Pattern completion via cell assembly retrieval was assessed with a metric we called pattern reconstruction. First, Pearson correlation coefficients (PCCs) were computed from the training and testing input and training and testing output as described previously [45]. Pattern reconstruction accuracy was then computed as the difference in the output and input PCCs divided by the difference of the maximum PCC (1) and the input PCC, multiplied by 100 to obtain a percentage (Supplementary Figure 2). Therefore, a non-zero pattern reconstruction accuracy would mean a cell assembly was retrieved, with 100% accuracy meaning perfect assembly retrieval.

### Overlap of cell assemblies

Associations between episodic memories in CA3 may be encoded by neurons shared between cell assemblies [21,22]. Based on the finding of an overlap of 4-5% being suitable for recall of individual and overlapping assemblies [21], and that pattern reconstruction accuracy at 97.5% pattern degradation was close to zero for assembly size 275 (1.68%; meaning that overlapping memories would not interfere with one another), an overlap of 5% was selected. Shared cells between each of three assemblies were randomly selected before training commenced, with the same training and synaptic downscaling procedures for no shared cells (0% overlap) utilized to obtain and normalize the weights between overlapping cell assemblies, respectively. For testing pattern completion with overlaps, an equal proportion of overlapping and non-overlapping cell assembly members were selected for stimulation, e.g., for an assembly size of 300 tested with a degraded pattern of 50%, 135 non-overlapping members and 15 overlapping members were randomly selected to activate each of the three assemblies.

### Model implementation and execution

The CA3 model was implemented in CARLsim6 [39], which utilized the 4^th^ order Runge Kutta numerical integration method with a fixed time step of 0.2 ms [72]. The duration for simulations that trained and tested the networks were 70 s and 1 s, respectively. Instantiation and execution of the network model was performed on single 40 and 80 GB VRAM Tesla A100 GPUs on the George Mason University High Performance Computing Cluster (Hopper). Hopper, which contained more than one hundred such GPUs, allowed for efficient and flexible simulation that greatly reduced the time needed to test different training and testing paradigms. Simulation results were loaded and visualized in MATLAB with CARLsim6’s Offline Analysis Toolbox (OAT). Additional custom-built functions for data analysis were written in Python and MATLAB. All scripts developed are available open source at github.com/jkopsick/cell_assembly_formation_retrieval.

## Acknowledgements

The authors are grateful to Drs. Carolina Tecuatl, Diek Wheeler, Rebecca Goldin, Nate Sutton, and Jonah Ascoli for helpful discussions.

## Author Contributions

Conceptualization: Jeffrey D. Kopsick, Giorgio A. Ascoli

Data curation: Jeffrey D. Kopsick, Joseph A. Kilgore

Formal analysis: Jeffrey D. Kopsick, Giorgio A. Ascoli

Funding Acquisition: Gina C. Adam, Giorgio A. Ascoli

Investigation: Jeffrey D. Kopsick, Giorgio A. Ascoli

Methodology: Jeffrey D. Kopsick, Giorgio A. Ascoli

Project Administration: Giorgio A. Ascoli

Resources: Gina C. Adam, Giorgio A. Ascoli

Software: Jeffrey D. Kopsick, Joseph A. Kilgore

Supervision: Gina C. Adam, Giorgio A. Ascoli

Validation: Jeffrey D. Kopsick, Giorgio A. Ascoli

Visualization: Jeffrey D. Kopsick

Writing – original draft: Jeffrey D. Kopsick

Writing – review & editing: Jeffrey D. Kopsick, Joseph A. Kilgore, Gina C. Adam, Giorgio A. Ascoli

## Funding

This work was supported by Department of Energy (DOE) grants DE-SC0022998 and DE-SC00023000, and National Institutes of Health (NIH) grant R01NS39600. The funders had no role in study design, data collection and analysis, decision to publish, or preparation of the manuscript.

## Data availability

All data used in this study is publicly available at https://doi.org/10.5281/zenodo.10870586.

## Notes

### Competing Interest Statement

The authors have declared no competing interest.

## References

1. Eichenbaum H. Hippocampus: Cognitive Processes and Neural Representations that Underlie Declarative Memory. Neuron. 2004 Sep 30;44(1):109–20.

2. Pfeiffer BE. The content of hippocampal “replay.” Hippocampus. 2020;30(1):6–18.

3. Dragoi G, Tonegawa S. Preplay of future place cell sequences by hippocampal cellular assemblies. Nature. 2011 Jan;469(7330):397–401.

4. Stachenfeld KL, Botvinick MM, Gershman SJ. The hippocampus as a predictive map. Nature Neuroscience. 2017 Nov;20(11):1643–53.

5. Rolls ET. The storage and recall of memories in the hippocampo-cortical system. Cell Tissue Res. 2018 Sep 1;373(3):577–604.

6. Rebola N, Carta M, Mulle C. Operation and plasticity of hippocampal CA3 circuits: implications for memory encoding. Nat Rev Neurosci. 2017 Apr;18(4):208–20.

7. Buzsáki G. Neural syntax: cell assemblies, synapsembles and readers. Neuron. 2010 Nov 4;68(3):362–85.

8. Farooq U, Sibille J, Liu K, Dragoi G. Strengthened Temporal Coordination within Pre-existing Sequential Cell Assemblies Supports Trajectory Replay. Neuron. 2019 Aug 21;103(4):719–733.e7.

9. Feldman DE. The spike timing dependence of plasticity. Neuron. 2012 Aug 23;75(4):556– 71.

10. Perez-Rosello T, Baker JL, Ferrante M, Iyengar S, Ascoli GA, Barrionuevo G. Passive and active shaping of unitary responses from associational/commissural and perforant path synapses in hippocampal CA3 pyramidal cells. J Comput Neurosci. 2011 Oct 1;31(2):159– 82.

11. Debanne D, Gähwiler BH, Thompson SM. Long-term synaptic plasticity between pairs of individual CA3 pyramidal cells in rat hippocampal slice cultures. J Physiol. 1998 Feb 15;507(Pt 1):237–47.

12. Mishra RK, Kim S, Guzman SJ, Jonas P. Symmetric spike timing-dependent plasticity at CA3– CA3 synapses optimizes storage and recall in autoassociative networks. Nat Commun. 2016 May 13;7(1):11552.

13. Kakegawa W, Tsuzuki K, Yoshida Y, Kameyama K, Ozawa S. Input- and subunit-specific AMPA receptor trafficking underlying long-term potentiation at hippocampal CA3 synapses. European Journal of Neuroscience. 2004;20(1):101–10.

14. Neunuebel JP, Knierim JJ. CA3 Retrieves Coherent Representations from Degraded Input: Direct Evidence for CA3 Pattern Completion and Dentate Gyrus Pattern Separation. Neuron. 2014 Jan 22;81(2):416–27.

15. Lazarewicz MT, Migliore M, Ascoli GA. A new bursting model of CA3 pyramidal cell physiology suggests multiple locations for spike initiation. Biosystems. 2002 Oct 1;67(1):129–37.

16. Hemond P, Epstein D, Boley A, Migliore M, Ascoli GA, Jaffe DB. Distinct classes of pyramidal cells exhibit mutually exclusive firing patterns in hippocampal area CA3b. Hippocampus. 2008;18(4):411–24.

17. Hemond P, Migliore M, Ascoli GA, Jaffe DB. The membrane response of hippocampal CA3b pyramidal neurons near rest: Heterogeneity of passive properties and the contribution of hyperpolarization-activated currents. Neuroscience. 2009 May 5;160(2):359–70.

18. Ascoli GA, Brown KM, Calixto E, Card JP, Galvan EJ, Perez-Rosello T, et al. Quantitative Morphometry of Electrophysiologically Identified CA3b Interneurons Reveals Robust Local Geometry and Distinct Cell Classes. J Comp Neurol. 2009 Aug 20;515(6):677–95.

19. de Almeida L, Idiart M, Lisman JE. Memory retrieval time and memory capacity of the CA3 network: Role of gamma frequency oscillations. Learn Mem. 2007 Nov;14(11):795–806.

20. Guzman SJ, Schlögl A, Frotscher M, Jonas P. Synaptic mechanisms of pattern completion in the hippocampal CA3 network. Science. 2016 Sep 9;353(6304):1117–23.

21. Gastaldi C, Schwalger T, Falco ED, Quiroga RQ, Gerstner W. When shared concept cells support associations: Theory of overlapping memory engrams. PLOS Computational Biology. 2021 Dec 30;17(12):e1009691.

22. Quian Quiroga R. An integrative view of human hippocampal function: Differences with other species and capacity considerations. Hippocampus. 2023;33(5):616–34.

23. Treves A, Rolls ET. What determines the capacity of autoassociative memories in the brain? Network. 1991 Nov;2(4):371.

24. Wheeler DW, White CM, Rees CL, Komendantov AO, Hamilton DJ, Ascoli GA. Hippocampome.org: a knowledge base of neuron types in the rodent hippocampus. elife. 2015 Sep 24;4.

25. Wheeler DW, Kopsick JD, Sutton N, Tecuatl C, Komendantov AO, Nadella K, et al. Hippocampome.org 2.0 is a knowledge base enabling data-driven spiking neural network simulations of rodent hippocampal circuits. Scharfman HE, Colgin LL, editors. eLife. 2024 Feb 12;12:RP90597.

26. White CM, Rees CL, Wheeler DW, Hamilton DJ, Ascoli GA. Molecular expression profiles of morphologically defined hippocampal neuron types: Empirical evidence and relational inferences. Hippocampus. 2020;30(5):472–87.

27. Ascoli GA, Wheeler DW. In search of a periodic table of the neurons: Axonal-dendritic circuitry as the organizing principle. BioEssays. 2016;38(10):969–76.

28. Komendantov AO, Venkadesh S, Rees CL, Wheeler DW, Hamilton DJ, Ascoli GA. Quantitative firing pattern phenotyping of hippocampal neuron types. Sci Rep. 2019 Nov 29;9.

29. Sanchez-Aguilera A, Wheeler DW, Jurado-Parras T, Valero M, Nokia MS, Cid E, et al. An update to Hippocampome.org by integrating single-cell phenotypes with circuit function in vivo. PLOS Biology. 2021 May 6;19(5):e3001213.

30. Attili SM, Silva MFM, Nguyen T vi, Ascoli GA. Cell numbers, distribution, shape, and regional variation throughout the murine hippocampal formation from the adult brain Allen Reference Atlas. Brain Struct Funct. 2019 Nov 1;224(8):2883–97.

31. Attili SM, Moradi K, Wheeler DW, Ascoli GA. Quantification of neuron types in the rodent hippocampal formation by data mining and numerical optimization. European Journal of Neuroscience. 2022;55(7):1724–41.

32. Rees CL, Moradi K, Ascoli GA. Weighing the Evidence in Peters’ Rule: Does Neuronal Morphology Predict Connectivity? Trends in Neurosciences. 2017;40(2):63–71.

33. Moradi K, Ascoli GA. Systematic Data Mining of Hippocampal Synaptic Properties. In: Cutsuridis V, Graham BP, Cobb S, Vida I, editors. Hippocampal Microcircuits: A Computational Modeler’s Resource Book. Springer International Publishing; 2018. p. 441–71. (Springer Series in Computational Neuroscience).

34. Moradi K, Ascoli GA. A comprehensive knowledge base of synaptic electrophysiology in the rodent hippocampal formation. Hippocampus. 2020;30(4):314–31.

35. Tecuatl C, Wheeler DW, Sutton N, Ascoli GA. Comprehensive Estimates of Potential Synaptic Connections in Local Circuits of the Rodent Hippocampal Formation by Axonal-Dendritic Overlap. J Neurosci. 2021 Feb 24;41(8):1665–83.

36. Venkadesh S, Komendantov AO, Wheeler DW, Hamilton DJ, Ascoli GA. Simple models of quantitative firing phenotypes in hippocampal neurons: Comprehensive coverage of intrinsic diversity. PLOS Computational Biology. 2019 Oct 28;15(10):e1007462.

37. Moradi K, Aldarraji Z, Luthra M, Madison GP, Ascoli GA. Normalized unitary synaptic signaling of the hippocampus and entorhinal cortex predicted by deep learning of experimental recordings. Commun Biol. 2022 May 5;5(1):1–19.

38. Kopsick JD, Tecuatl C, Moradi K, Attili SM, Kashyap HJ, Xing J, et al. Robust Resting-State Dynamics in a Large-Scale Spiking Neural Network Model of Area CA3 in the Mouse Hippocampus. Cogn Comput. 2023 Jul 1;15(4):1190–210.

39. Niedermeier L, Chen K, Xing J, Das A, Kopsick JD, Scott EO, et al. CARLsim 6: An Open Source Library for Large-Scale, Biologically Detailed Spiking Neural Network Simulation. In: 2022 International Joint Conference on Neural Networks (IJCNN).

40. Lisman JE, Jensen O. The Theta-Gamma Neural Code. Neuron. 2013 Mar 20;77(6):1002–16.

41. Tononi G, Cirelli C. Sleep and synaptic homeostasis: a hypothesis. Brain Research Bulletin. 2003 Dec;62(2):143–50.

42. Tononi G, Cirelli C. Sleep and the Price of Plasticity: From Synaptic and Cellular Homeostasis to Memory Consolidation and Integration. Neuron. 2014 Jan;81(1):12–34.

43. Tononi G, Cirelli C. Sleep function and synaptic homeostasis. Sleep Medicine Reviews. 2006 Feb;10(1):49–62.

44. Hebb DO. The Organization of Behavior. John Wiley & Sons; 1949.

45. Guzman SJ, Schlögl A, Espinoza C, Zhang X, Suter BA, Jonas P. How connectivity rules and synaptic properties shape the efficacy of pattern separation in the entorhinal cortex– dentate gyrus–CA3 network. Nat Comput Sci. 2021 Dec;1(12):830–42.

46. Waydo S, Kraskov A, Quiroga RQ, Fried I, Koch C. Sparse Representation in the Human Medial Temporal Lobe. J Neurosci. 2006 Oct 4;26(40):10232–4.

47. Bennett MR, Gibson WG, Robinson J. Dynamics of the CA3 pyramidial neuron autoassociative memory network in the hippocampus. Philosophical Transactions of the Royal Society of London Series B: Biological Sciences. 1997 Jan;343(1304):167–87.

48. Lisman JE. Relating Hippocampal Circuitry to Function: Recall of Memory Sequences by Reciprocal Dentate–CA3 Interactions. Neuron. 1999 Feb 1;22(2):233–42.

49. Kemker R, McClure M, Abitino A, Hayes T, Kanan C. Measuring Catastrophic Forgetting in Neural Networks. Proceedings of the AAAI Conference on Artificial Intelligence. 2018 Apr 29;32(1).

50. Kumaran D, Hassabis D, McClelland JL. What Learning Systems do Intelligent Agents Need? Complementary Learning Systems Theory Updated. Trends in Cognitive Sciences. 2016 Jul 1;20(7):512–34.

51. Moser EI, Moser MB. One-Shot Memory in Hippocampal CA3 Networks. Neuron. 2003 Apr 24;38(2):147–8.

52. Nakazawa K, Quirk MC, Chitwood RA, Watanabe M, Yeckel MF, Sun LD, et al. Requirement for Hippocampal CA3 NMDA Receptors in Associative Memory Recall. Science. 2002 Jul 12;297(5579):211–8.

53. Montgomery SM, Buzsáki G. Gamma oscillations dynamically couple hippocampal CA3 and CA1 regions during memory task performance. Proc Natl Acad Sci U S A. 2007 Sep 4;104(36):14495–500.

54. Neves L, Lobão-Soares B, Araujo AP de C, Furtunato AMB, Paiva I, Souza N, et al. Theta and gamma oscillations in the rat hippocampus support the discrimination of object displacement in a recognition memory task. Front Behav Neurosci. 2022 Dec 21;16:970083.

55. Siegle JH, Wilson MA. Enhancement of encoding and retrieval functions through theta phase-specific manipulation of hippocampus. Eichenbaum H, editor. eLife. 2014 Jul 29;3:e03061.

56. Buzsáki G. The Brain from Inside Out. New York, NY: Oxford University Press; 2019. 464 p.

57. González-Rueda A, Pedrosa V, Feord RC, Clopath C, Paulsen O. Activity-Dependent Downscaling of Subthreshold Synaptic Inputs during Slow-Wave-Sleep-like Activity In Vivo. Neuron. 2018 Mar 21;97(6):1244–1252.e5.

58. Hummos A, Franklin CC, Nair SS. Intrinsic mechanisms stabilize encoding and retrieval circuits differentially in a hippocampal network model. Hippocampus. 2014;24(12):1430– 48.

59. Sammons RP, Vezir M, Moreno-Velasquez L, Cano G, Orlando M, Sievers M, et al. Structure and function of the hippocampal CA3 module. Proceedings of the National Academy of Sciences. 2024 Feb 6;121(6):e2312281120.

60. Carrillo-Reid L, Han S, Yang W, Akrouh A, Yuste R. Controlling Visually Guided Behavior by Holographic Recalling of Cortical Ensembles. Cell. 2019 Jul 11;178(2):447–457.e5.

61. Quiroga RQ. Concept cells: the building blocks of declarative memory functions. Nat Rev Neurosci. 2012 Aug;13(8):587–97.

62. Steinmetz NA, Aydin C, Lebedeva A, Okun M, Pachitariu M, Bauza M, et al. Neuropixels 2.0: A miniaturized high-density probe for stable, long-term brain recordings. Science. 2021 Apr 16;372(6539):eabf4588.

63. Zong W, Obenhaus HA, Skytøen ER, Eneqvist H, de Jong NL, Vale R, et al. Large-scale two-photon calcium imaging in freely moving mice. Cell. 2022 Mar 31;185(7):1240–1256.e30.

64. Vöröslakos M, Kim K, Slager N, Ko E, Oh S, Parizi SS, et al. HectoSTAR μLED Optoelectrodes for Large-Scale, High-Precision In Vivo Opto-Electrophysiology. Advanced Science. 2022 Apr 22;n/a(n/a):2105414.

65. Knierim JJ. Dynamic Interactions between Local Surface Cues, Distal Landmarks, and Intrinsic Circuitry in Hippocampal Place Cells. J Neurosci. 2002 Jul 15;22(14):6254–64.

66. Izhikevich EM. Dynamical Systems in Neuroscience. MIT Press; 2007. 522 p.

67. Tsodyks M, Pawelzik K, Markram H. Neural Networks with Dynamic Synapses. Neural Computation. 1998 May 15;10(4):821–35.

68. Tecuatl C, Wheeler DW, Ascoli GA. A Method for Estimating the Potential Synaptic Connections Between Axons and Dendrites From 2D Neuronal Images. Bio-protocol. 2021 Jul 5;11(13):e4073–e4073.

69. Mizuseki K, Buzsáki G. Preconfigured, Skewed Distribution of Firing Rates in the Hippocampus and Entorhinal Cortex. Cell Reports. 2013 Sep 12;4(5):1010–21.

70. Buzsáki G, Mizuseki K. The log-dynamic brain: how skewed distributions affect network operations. Nat Rev Neurosci. 2014 Apr;15(4):264–78.

71. Miles R, Le Duigou C, Simonnet J, Telenczuk M, Fricker D. Recurrent synapses and circuits in the CA3 region of the hippocampus: an associative network. Frontiers in Cellular Neuroscience. 2014;7.

72. Butcher JC. A history of Runge-Kutta methods. Applied Numerical Mathematics. 1996 Mar 1;20(3):247–60.

73. Kay K, Sosa M, Chung JE, Karlsson MP, Larkin MC, Frank LM. A hippocampal network for spatial coding during immobility and sleep. Nature. 2016 Mar;531(7593):185–90.

74. Lasztóczi B, Tukker JJ, Somogyi P, Klausberger T. Terminal Field and Firing Selectivity of Cholecystokinin-Expressing Interneurons in the Hippocampal CA3 Area. J Neurosci. 2011 Dec 7;31(49):18073–93.

75. Oliva A, Fernández-Ruiz A, Buzsáki G, Berényi A. Spatial coding and physiological properties of hippocampal neurons in the Cornu Ammonis subregions. Hippocampus. 2016;26(12):1593–607.

76. Ding L, Chen H, Diamantaki M, Coletta S, Preston-Ferrer P, Burgalossi A. Structural Correlates of CA2 and CA3 Pyramidal Cell Activity in Freely-Moving Mice. J Neurosci. 2020 Jul 22;40(30):5797–806.

77. Viney TJ, Lasztoczi B, Katona L, Crump MG, Tukker JJ, Klausberger T, et al. Network state-dependent inhibition of identified hippocampal CA3 axo-axonic cells in vivo. Nature Neuroscience. 2013 Dec;16(12):1802–11.

78. Tukker JJ, Lasztóczi B, Katona L, Roberts JDB, Pissadaki EK, Dalezios Y, et al. Distinct Dendritic Arborization and In Vivo Firing Patterns of Parvalbumin-Expressing Basket Cells in the Hippocampal Area CA3. J Neurosci. 2013 Apr 17;33(16):6809–25.

79. Lapray D, Lasztoczi B, Lagler M, Viney TJ, Katona L, Valenti O, et al. Behavior-dependent specialization of identified hippocampal interneurons. Nature Neuroscience. 2012 Sep;15(9):1265–71.

80. Varga C, Golshani P, Soltesz I. Frequency-invariant temporal ordering of interneuronal discharges during hippocampal oscillations in awake mice. PNAS. 2012 Oct 2;109(40):E2726–34.

81. Klausberger T, Márton LF, Baude A, Roberts JDB, Magill PJ, Somogyi P. Spike timing of dendrite-targeting bistratified cells during hippocampal network oscillations in vivo. Nature Neuroscience. 2004 Jan;7(1):41–7.

82. Katona L, Lapray D, Viney TJ, Oulhaj A, Borhegyi Z, Micklem BR, et al. Sleep and Movement Differentiates Actions of Two Types of Somatostatin-Expressing GABAergic Interneuron in Rat Hippocampus. Neuron. 2014 May 21;82(4):872–86.

83. Fuentealba P, Begum R, Capogna M, Jinno S, Márton LF, Csicsvari J, et al. Ivy Cells: A Population of Nitric-Oxide-Producing, Slow-Spiking GABAergic Neurons and Their Involvement in Hippocampal Network Activity. Neuron. 2008 Mar 27;57(6):917–29.

